# Label-free 3D virtual histology of human formalin-fixed paraffin-embedded (FFPE) prostate needle biopsies with propagation-based phase-contrast micro-CT (PBCT)

**DOI:** 10.64898/2026.05.28.728215

**Authors:** Andrew Sugarman, Daniel Vanselow, Guoli Chen, Elizabeth Clark, Dilworth Parkinson, Patrick La Riviere, Justin Silverman, Joshua Warrick, Keith C. Cheng

## Abstract

For over a century, the goal of estimating clinical outcome from tumor biopsies has been based on histomorphology of 2D tissue slices that represent a small fraction of collected samples. Its power derives from histology’s 1) *unbiased representation* of cell types, 2) *subcellular resolution* that allows the characterization of health and disease states across cell types, and 3) *multi*-*millimeter fields of view* that allow assessment of tumor heterogeneity. Histology’s dependence upon physical slices, however, limits assessment of 3-dimensional cellular volumes and tissue architecture. Here, we used propagation-based phase-contrast micro-CT (PBCT) to create 3D histological images of residual formalin-fixed, paraffin-embedded (FFPE) prostate needle biopsies. The resulting isotropic, grey-scale, 0.5 micron voxel matrices were used to explore the potential of for the 3D virtual histology to distinguish diagnostic categories including benign prostatic tissue and prostatic adenocarcinoma of Gleason patterns 3, 4, and 5. Maximum intensity projections of stacks of digital slices totaling 5 microns “slices” allowed the study of virtual sections corresponding to actual serial H&E-stained sections of tissue cut after micro-CT imaging. Like histology, our PBCT reconstructions allowed us to distinguish between non-infiltrative and undulating glands of benign prostatic tissue, infiltrative round glands of Gleason pattern 3, cribriform structures of Gleason pattern 4, and comedonecrosis of Gleason pattern 5. Unlike histology, micro-CT allowed us to further probe 3D tissue architecture in volumetric context. User-friendly exploration of sample volumes was achieved using a customized Neuroglancer multiplanar and 3D rendering interface. Sparsely trained cycleGAN produced plausible virtual H&E staining from the unstained micro-CT reconstructions. Unlike tissue-section based histology, micro-CT-based virtual histology yields nondestructive 3D characterization of cancer cell and tissue architecture, including glandular spaces, without the undersampling or cutting artifacts of histology. These findings demonstrate the feasibility of PBCT-based 3D virtual histology of prostate cancer and suggest the exploration of derived quantitative analyses of tumor properties for potential contributions to patient care.

---

Prostate cancer (PCa) is the most common cancer in United States in men and a leading cause of cancer-related death worldwide (Kratzer et al., 2025; Siegel et al., 2024). Tumors differ dramatically in aggressiveness and clinical outcomes and can show tremendous clonal heterogeneity (Haffner et al., 2015). Standard diagnosis of PCa relies on the qualitative assessment of 2D histological sections taken from transrectal or transperineal needle core biopsies. Hematoxylin and Eosin-stained (H&E) slides from each needle core are then assigned a Gleason score by combining their 2 most prevalent Gleason patterns, a summary statistic for tumor severity based on the architecture of visible prostate glands (Rebello et al., 2021). To make treatment decisions in clinical practice, patients are then stratified into an ISUP (International Society of Urological Pathology) grade group derived from the Gleason grade measured via histology of needle-core biopsies. Individuals with Grade Group 1 and some with Grade Group 2 tumors are candidates for active surveillance. Those with Grade Group 5 are treated aggressively. Gleason score is the single most important determinant of treatment. Limitations of Gleason grading include under sampling both in the gathering of needle biopsy material, percent inspection of the biopsy sample using standard H&E sections, and limited volumetric assessment of cells and cellular architecture. Risk stratification within grade groups is an ongoing challenge (Gordetsky & Epstein, 2016; van Leenders et al., 2020).

High-resolution 3D imaging is a potential way to address the challenge of assaying sampled tissues volumetrically. Work has begun towards quantifying the 3D structures that determine qualitative 2D phenotypes, such as gland shape using fluorescence microscopy (Wang et al., 2024). Fluorescence-based open-top light-sheet microscopy (OTLS) exemplifies the potential translational benefit of volumetric imaging, particularly in the prediction of biochemical recurrence (Glaser et al., 2022; Liu et al., 2021; Reder et al., 2019; Serafin et al., 2023; Song et al., 2024; Wang et al., 2024; Xie et al., 2022). These approaches require deparaffinization for embedded tissues, tissue clearing, and fluorescent staining, which lie outside the conventional workflow of histopathology (Daetwyler & Fiolka, 2023; Glaser et al., 2022; Liu et al., 2021).

Phase-contrast X-ray microscopy is able to resolve soft tissue details of low refractive index without the addition of contrast enhancing stains commonly used for micro-CT to maximize signal in biological samples (Arana Peña et al., 2023; Frohn et al., 2020; Pinkert-Leetsch et al., 2023; Topperwien et al., 2019; Töpperwien et al., 2018, 2020). Recent advances in micro-CT tailored for soft tissue - referred to as histotomography – have extended sub-micrometer resolution to samples to 10 millimeters (Ding et al., 2019; Ngu et al., 2024; Xin et al., 2017; Yakovlev et al., 2022, 2023). Histotomography, performed with or without metal staining, utilizes differences in physical density and pattern between cellular components such as nucleic acid and structural proteins, resulting in spatially resolved morphological patterns. The characteristics that have made it a useful form of 3D histology include isotropic sub-micrometer resolution, pan-cellular detection based on different densities of cellular and extracellular tissue components as well as phase-based edge detection. Proof-of-principle of 3D histology using micro-CT was established using phosphotungstic acid (PTA) or silver staining to boost signal (Ding et al., 2019; Katz et al., 2021; Ngu et al., 2024; Yakovlev et al., 2022), but here we take advantage of knowledge that density differences present without metal stain result in sufficient phase effects to create remarkable similarity to histological images. We focus here on the potential of propagation-based phase-contrast micro-CT (PBCT) because of its minimal interference with prevailing handling of tumor tissue. PBCT offers isotropic sub-micrometer resolution, pan-cellular detection arising from density differences between cellular and extracellular tissue components, and phase-based edge enhancement – without requiring deparaffinization or contrast-enhancing stains.

In this study, we test the potential of the extension of histotomography to leverage propagation-based phase-contrast for volumetric imaging of unstained, formalin-fixed paraffin embedded (FFPE) prostate needle biopsies. Micro-CT images were directly compared with standard H&E-stained sections in both benign and malignant examples. PBCT allowed qualitative description of the full imaged volume of prostate tissue biopsies including morphological assessments through the full sample, setting a potentially useful stage for quantitative morphometric analytics.

## Materials and Methods

### Sample selection and preparation

Human prostate needle-core biopsies were selected from a cohort of patients based on their Gleason score at initial diagnosis. Each patient in this study eventually went on to have a radical prostatectomy, but tissue blocks were identified to contain major phenotypes including benign tissue, Gleason pattern 3, 4, and 5. Four needle biopsies were chosen for comparative micro-CT and H&E examination: one benign prostatic tissue, one Gleason score 3+3=6 (Grade Group 1), one Gleason score 4+4=8 (Grade Group 4) with cribriform histology, and one Gleason score 4+5=9 (Grade Group 5) with comedonecrosis (Table S1). Archived H&E slides and FFPE blocks from prostate needle-core biopsies were identified through PathNet and obtained with IRB approval (STUDY00023526) from the Pathology Department at Hershey Medical Center (HMC). Gleason scores for the present study were based on review by study pathologists (GS, JW). For micro-CT imaging, remnant core biopsy was utilized, left embedded in paraffin for scanning, though excess paraffin was manually removed with a scalpel for micro-CT scanning. In the case where biopsies were folded or embedded in pieces, one representative piece was extracted from the tissue block with a scalpel. Individual paraffin-embedded biopsy tissues were temporarily affixed to plastic sample holders using excess paraffin wax.

### Synchrotron micro-CT imaging

Synchrotron propagation-based phase contrast micro-CT (PBCT) soft tissue imaging utilizes refraction and diffraction of X-rays by component substances to produce enhanced contrast between component material of different density, from dense nuclei to less dense cytoplasmic features. As X-rays pass through the sample, phase changes are induced by materials of varying electron densities, causing interference as the wave propagates from the sample to the detector, where they manifest as variations in image intensity. In mathematical terms, induced intensity distribution of a PBCT image is governed by the sample’s complex index of refraction (n), which in turn is a combination of absorption (β) and phase effects (δ):

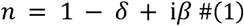

For weakly-absorbing samples with refractive indices around 1, the phase component (*δ*) of this equation is thought to be primarily responsible for captured intensity (Jacobsen, 2019; Paganin, 2006; Paganin & Pelliccia, 2021; Topperwien et al., 2019; Töpperwien et al., 2018). The clarity of an image generated by PBCT, ie the magnitude of refraction and the diffraction patterns observed in the plane of the detector, is governed by the Fresnel number (see Eq. 2). The Fresnel number is dependent upon feature size *a*, beam wavelength *λ*, and propagation distance *R* (Equation 2, Figure 1A).

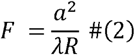

The goal of the presented experiments was to maximize contrast enhancement while minimizing distorting artifact and blur. We chose to image within the direct-contrast regime (*F* ∼ 1) for cell nuclei of roughly 5*µ*m in diameter, with the goal of enhancing material edges while not introducing multiple “fringes” within the reconstruction (*F* ≪ 1, holographic regime). At beamline 8.3.2, we selected a beam energy of 14 keV due to its relatively high 2-3 mm vertical dimension and for an increased contribution of photon absorption (β) to our images compared with lower contrast obtained at higher energies. We empirically tested a series of propagation distances with 14 keV X-rays (Figure. S1), ultimately choosing *R* = 50 mm as an optimal balance between contrast and perceptible blur. Use of 70 mm propagation distance at 14 keV, for example, yielded strong contrast, but with increased blur. We calculated roughly *F* = 1.41 for a feature size of nuclear radius (∼2.5 um), 14 keV (λ = 0.08856 nanometers), and *R* = 50 mm (Beltran et al., 2010; Jacobsen, 2019). This Fresnel number aligns with the direct contrast regime (*F*≃1), such that fringes will appear as edge enhancement between materials of different density within the sample. By performing PBCT within the direct-contrast regime, we were able to utilize the Bronnikov-Aided Correction (BAC) for single-distance phase retrieval, which does not require the single-material approximation (Lohse et al., 2020; Witte et al., 2009). All synchrotron imaging was performed at beamline 8.3.2 of Lawrence Berkeley National Laboratory’s Advanced Light Source (ALS) over the course of 2 beam time allocations.

**Figure 1:**
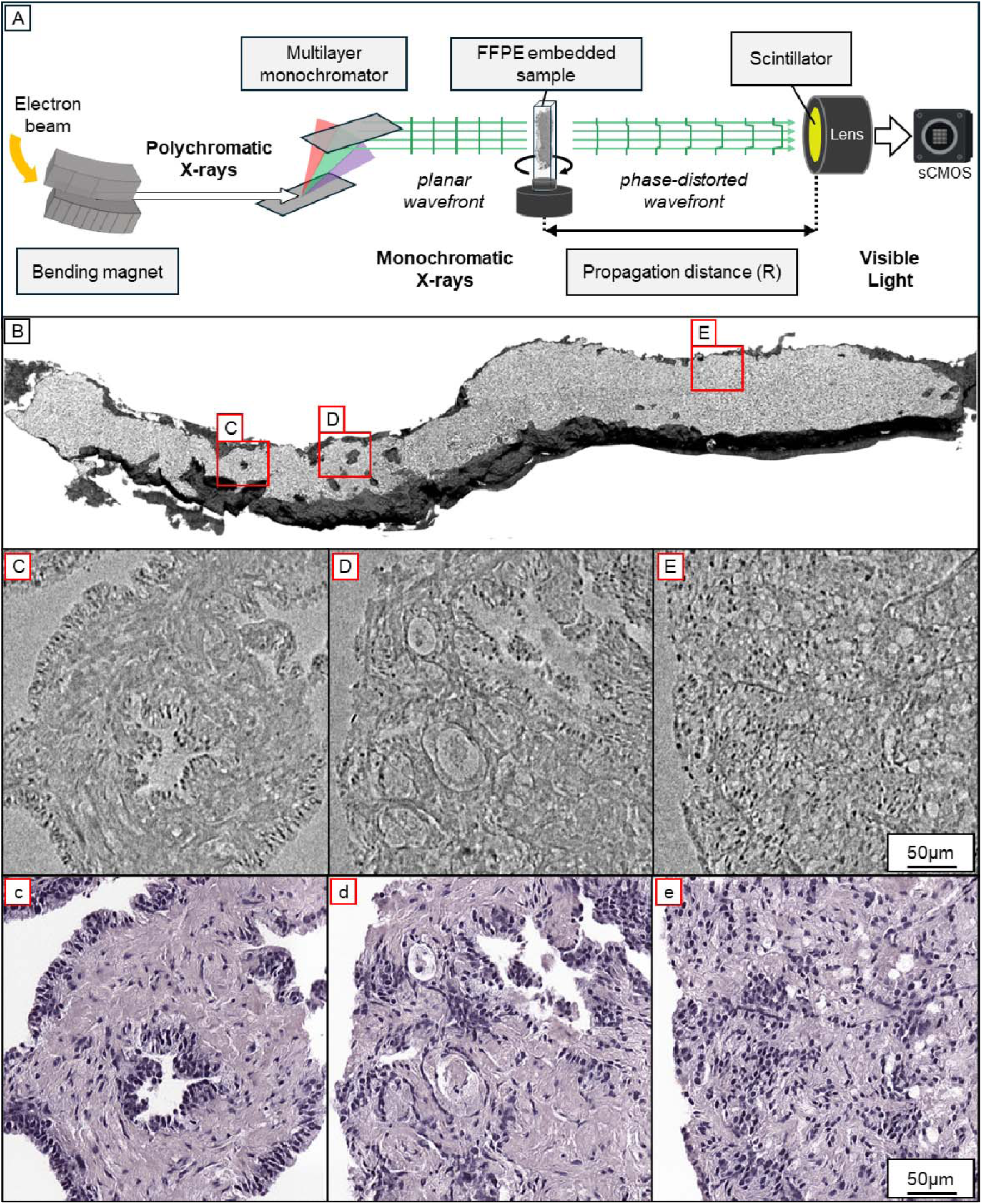
Propagation-based phase contrast micro-CT (PBCT) enables visualization of prostate glands and nuclei without addition of contrast-enhancing stain. (A) Schematic of the parallel-beam synchrotron micro-CT setup at Lawrence Berkeley National Laboratory. Conventional X-ray and CT imaging generate contrast from differences in how strongly tissues absorb X-rays; PBCT primarily detects subtle phase shifts imparted to the X-ray wavefront as it passes through tissue of differing electron density. These shifts produce no visible signal at the sample itself, but over a short free-space propagation distance (R) between the sample and the scintillator they develop into intensity differences that are strongest at the boundaries between components: the gland edges, stromal interfaces, and nuclear borders that define histologic morphology. Detection is indirect; the scintillator converts the transmitted X-rays to visible light, which the objective lens then projects onto the sCMOS camera that records the projection image. Lengthening R enhances this boundary contrast, allowing unstained formalin-fixed paraffin-embedded (FFPE) tissue to be imaged without any added stain. (B) 3D volumetric rendering of a biopsy core graded as Gleason pattern 4+4 (Case 3), captured with PBCT at 14keV and 50mm propagation distance. Red boxes indicate regions where c-e were extracted, with B showing a different depth chosen to highlight 3D features. (C-E) Single-slice (0.61 μm voxel size) regions of interest (ROIs) cropped from separate areas of the reconstruction rendered in (B) illustrate the variation in nuclear-scale morphology captured by PBCT. (C) Benign acini and fibromuscular stroma, (D) Mixed benign and abnormal acini, and (E) fused cribriform pattern 4 tumors are identifiable within the same reconstruction. (c-e) Panels C-E were virtually stained using a sparsely-trained cycleGAN to aid interpretation by those accustomed to H&E.

The four needle biopsies were scanned through a general user proposal (GUP) allocation whereby a custom wide-field micro-CT detector (described in Yakovlev, 2022) was transported to the imaging station at Lawrence Berkeley National Laboratory beamline 8.3.2 (Yakovlev et al., 2022). In preparation for synchrotron micro-CT imaging, excess paraffin (greater than roughly 0.5mm from the needle biopsy margin) was removed using heated 9, 10, or 11 scalpel blades to remove unnecessary material between the beam and the sample, enabling fixation to the sample rotation chuck, and to enable adjustment of sample-to-scintillator distance. Neither tissue clearing nor contrast-enhancing stains are necessary. Paraffin from the tissue block was used to fix the samples to the wide end of plastic needle caps (from Becton Dickenson Embecta 32 G Nano pen needles) and the narrow end of the caps were used for mounting in the sample holder for scanning. The full height of the samples were scanned with 2-7 tomograms per specimen using the custom 5mm detector or the 10x objective lens native to beamline 8.3.2. 1969 projections per tomogram were acquired and used to reconstruct the 3D tomograms with voxel sizes of 0.61 *µ*m and 0.65 *µ*m respectively.

For scans using the 5 mm detector system (Cases 1-4), 10-15 dark field then 10-15 flat field images were acquired before each tomogram (Yakovlev et al., 2022). 1969 projections were acquired with 300 ms exposure time in mono16 (16-bit) continuous scan mode across 180 degrees of sample rotation (approximately 10 minutes per tomogram). 10-15 post-scan flat field images were acquired after each tomogram for optimal angle correction during reconstruction. 10-15 still projections were also acquired at the starting angular position (0 degrees of rotation) to assay for sample movement during scanning. The angular deviation of our custom detector from the sample axis of rotation was calculated using a QRM phantom and by performing the center-finding step of reconstruction at both the top and the bottom of a scan, measuring the difference. This angle was found to be approximately 0.149, and projections were rotated to adjust for this in final reconstruction. For scans using LBNL’s native 0.65*µ*m voxel detector system at beamline 8.3.2, 1969 projections were acquired with an exposure time of 200 ms and no rotation correction was needed. For these scans, 50 mm of propagation distance and 14 keV beam energy were also used.

### Reconstruction, phase retrieval, and image stitching

Reconstruction was performed with a custom implementation of the TomoPy GridRec algorithm (Gursoy, 2014) in Python (Gürsoy et al., 2014). Center finding was performed through a combination of the PC center finding algorithm and manual assessment down to 0.25 micrometer step size for each sample. Phase retrieval was performed with the Bronnikov Aided Correction algorithm (De Witte, 2009) using parameter values alpha = 2 and gamma = 1.25 (Witte et al., 2009). BAC filtering was performed using the python implementation of the Holotomotoolbox named HoToPy (Lohse et al., 2020). Parameter values were ascertained through qualitative comparison of reconstructions processed with and without Bronnikov-aided correction (Figure S2). Reconstructed tomograms were cropped and aligned for stitching as needed using the Pairwise Stitcher plugin in ImageJ (Preibisch et al., 2009; Schindelin et al., 2012, 2015). Sections were fused using either the pairwise stitcher plugin with linear blending or with a custom python script using Skimage (see data and code availability).

### Correlative histology

The four needle biopsies (Table S1) were subjected to serial histological sectioning after micro-CT scanning. Biopsies were placed in a mold with the originally sectioned edge facing down. New paraffin was added to this mold to quickly reform a complete tissue block around the needle-core. The block was allowed to cool and new sections were cut at 5 *µ*m thickness until tissue was exhausted. These slides were stained with hematoxylin and eosin and affixed to slides for visualization by light microscope. Histology slide preparation was performed as follows: Every tissue section was baked at 37-40 degrees Celsius for 2 days to adhere them to glass slides. Slides were then baked at 60 degrees Celsius for one hour, then deparaffinized in xylene for 3 minutes 3 times. A serial ethanol wash was performed with 3 minutes in 100% ethanol 3 times, 3 minutes at 95% ethanol 3 times, and 70% for 3 minutes once. Slides were then rinsed in water for 3 minutes. Slides were stained with hematoxylin (Gill 3 from Avantik, #:11-5103-24) for 4.5 minutes, then rinsed with running tap water for 1 minute. This was followed by 4 dips with acid alcohol and another running tap water rinse for 1 minute. Slides were then treated with ammonia water for 30 seconds before another 1 minute of running tap water rinse and one minute of 95% ethanol. Slides were stained with eosin for 3 minutes, rinsed with 95% ethanol for 3 minutes 3 more times, then 100% ethanol for 3 minutes 3 more times. Slides were then treated with xylene for 3 minutes 3 times each and cover slipped. All histology slides cut after PBCT were scanned electronically using the Penn State College of Medicine, Department of Pathology Aperio slide scanner with 0.249 *µ*m micron effective pixel size.

### Segmentation and image processing

Five 256×256×256-voxel volumes of interest were cropped from the reconstruction of our 4+4 sample and likely nuclei were labeled with a LabKit classifier using hand-drawn annotations that were used to train and test a StarDist 3D U-Net classifier (Arzt et al., 2022; Weigert & Schmidt, 2022). Tiled inference using an A5000 GPU was performed on each section scan to yield 3D instance masks of candidate nuclei. Digital histology slide images generated by the Aperio scanner were post-processed with OpenCV in Python, and individual tissue pieces from each slide were automatically cropped and saved as individual images. Segmentations with the stock 2D StarDist algorithm in python yielded instance masks of nuclei.

Virtual hematoxylin and eosin staining was performed using a cycleGAN. 3 2D slices from micro-CT reconstructions were selected across cases 1 and 3 for training. Conventional H&E slides matching these representative tissue regions were also selected. We retrained our StarDist model with 12 256×256×256 voxel volumes from Cases 1-4 and performed nuclear segmentation for each image pair to facilitate further image registration. Nuclear masks were converted to density grids and further aligned using conventional deformable registration in skimage. A cycleGAN with a ResNet-9 generator and PatchGAN discriminator with least-squares adversarial loss was trained on 256×256 pixel image patches sampled from 512×512 pixel tiles. Tiles were partitioned into training, validation, and testing sets ahead of patch sampling. We used L1 cycle-consistency loss for the generator and set the identity term to 0 as the grayscale images were single channel and the H&E images were RGB. Ground-truth H&E images were stain-normalized using the Macenko procedure to ensure consistency (Macenko et al., 2009). Patches were sampled from 512×512 tiles generated from the 3 registered ROIs, such that tiles could be split into train/validation/testing sets ahead of sampling. 2000 patches were sampled per epoch, and 200 epochs were used for training. Checkpoints were saved every epoch. Inference was performed on 512-pixel tiles, which were stitched with 64-pixel overlap for larger test images.

## Results

### Phase-based micro-CT resolves morphological details of unstained prostate cancer in 3D

The density-based contrast offered by PBCT yielded digitally generated virtual “slices” of submicron thickness that closely resemble physical hematoxylin and eosin-stained tissue sections (Figures 2-6, Table S2). Like histology, the reconstructed images of our unstained biopsy cores revealed discernable nuclei due to greater X-ray absorption compared to cytoplasm, stroma and lumen. To increase resemblance of the images to histology, we inverted the reconstructed images so that nuclei appear dark, similar to appearance of hematoxylin and eosin-stained tissue histology.

**Figure 2:**
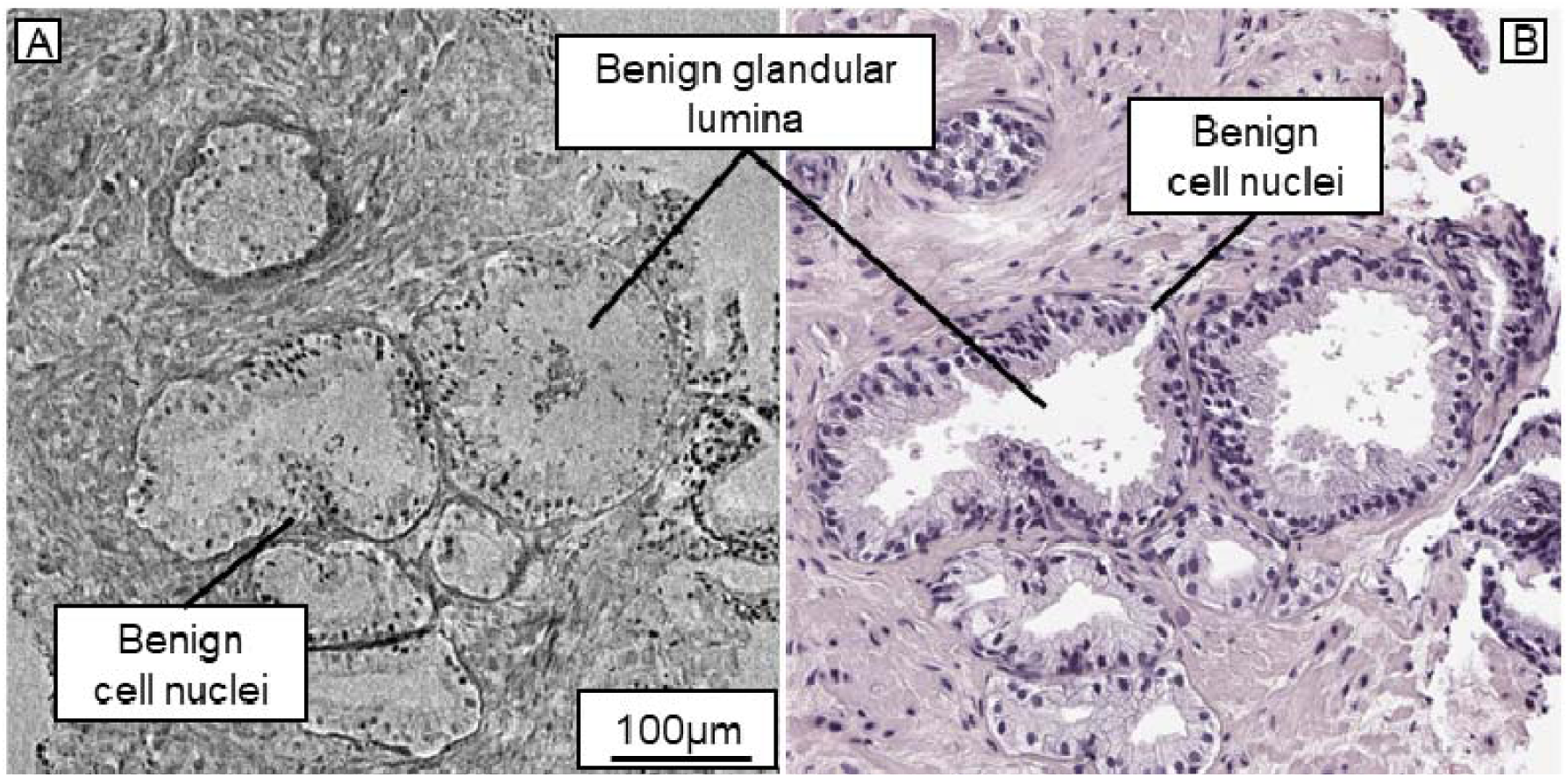
Features of benign prostatic tissue are visible across PBCT and correlative H&E sections. (A) 8-slice maximum intensity projection (∼4.8 micrometers thick) of a benign ROI cropped from the PBCT reconstruction of sample 1. (B) H&E-stained ROI from a section of the same tissue as (A), extracted after PBCT scanning. Glandular and nuclear patterns are discernible across imaging modalities.

Benign and malignant glands were distinguished based on luminal tortuosity, patterns of cell nuclei, and spatial arrangements of glands to one another (Figures 1-5). 3D images were resliced to test our ability to qualitatively represent different planes of histologic sectioning, and more importantly, to create orthogonal planes that enable alternate views of gland and cellular morphology beyond histology’s limited planes of section (Figure 3). The isotropic voxels of micro-CT enable virtual re-slicing in *any* plane, allowing 360-degree views throughout volumes of interest (Figure S3). We found that the major Gleason patterns used in the diagnosis and grading of prostate cancer are readily distinguishable from the reconstructed PBCT images (Figure S4). The combination of nuclear contrast and the visibility of lumen within PBCT images enabled fine discrimination of morphology across the range of magnifications of PBCT images Orthogonal virtual sections also revealed subsurface variation in glandular architecture beneath the original plane of sectioning (Figure S5). 2D grayscale slices were inspected in collaboration with 2 board-certified genitourinary pathologists (JW, GC) and compared with H&E slides from before and after PBCT scanning. Well-formed lumen and organized nuclear patterns constituting benign acini are resolved by PBCT and validated by matching H&E (Figures 2-6). To accurately represent the thickness of histopathology sections, we generated 5 *µ*m maximum intensity projections (MIPs), choosing digital planes of section that closely matched that of the physical sections. Notably, the simulated H&E sections produced are not marred by sectioning artifact such as folding artifact (seen in Figure 3).

**Figure 3:**
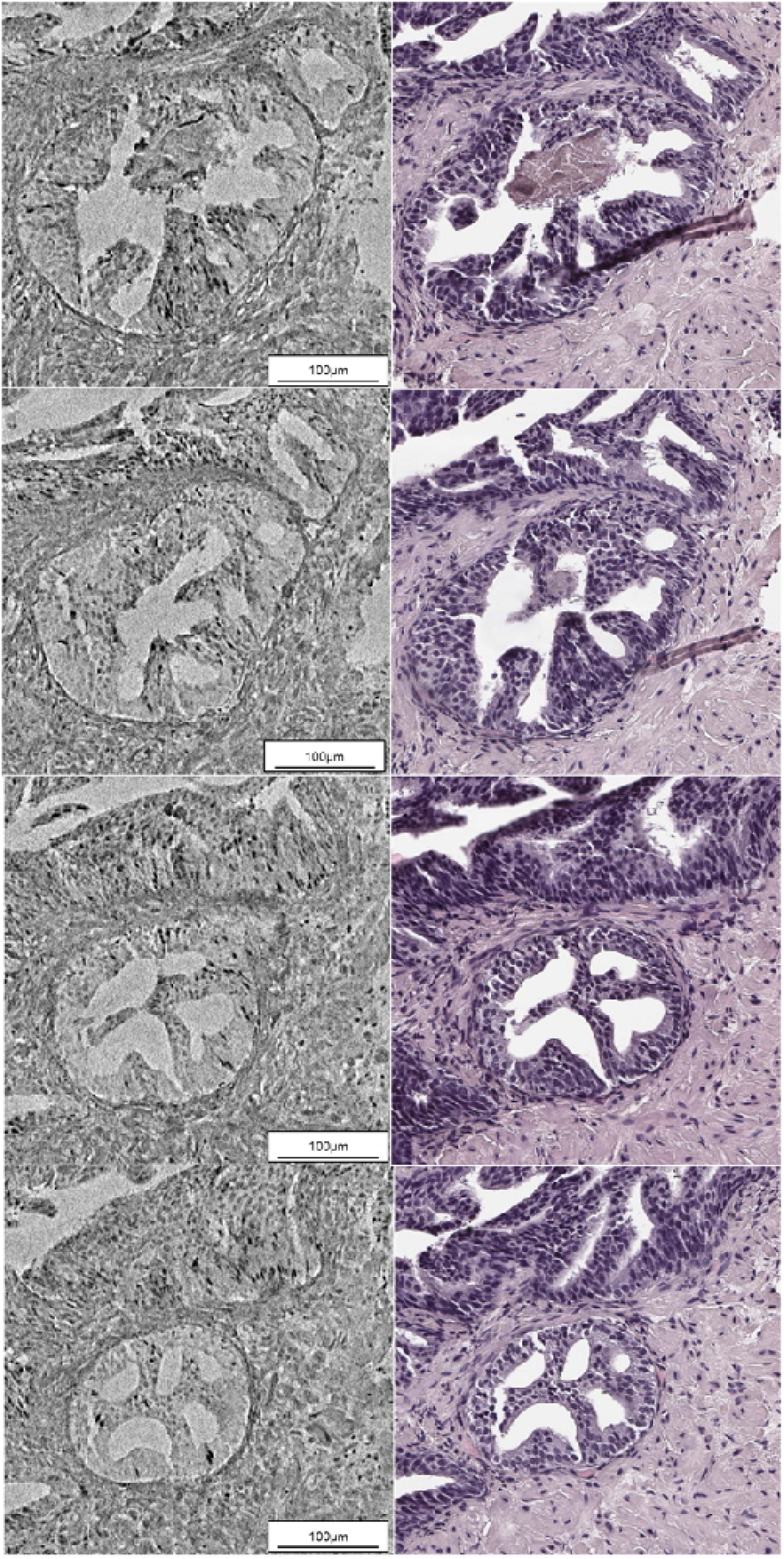
Variation in luminal microstructure revealed by depth-based interrogation of PBCT reconstruction with correlative histology. (A) ROI snapshots virtually sectioned from a PBCT reconstruction of a biopsy initially diagnosed as benign. Inverted 8-slice maximum intensity projections (∼4.8 micrometers of tissue) are shown to better match the contrast of histology. These snapshots were selected to qualitatively correspond to the plane of sectioning corresponding to initial diagnostic cuts and those performed for (B). Nuclei are visible as dark foci while gland lumina appear as light grey. MIPs in (A) are separated by 42 slices (∼25 micrometers) and shown from superficial (top) to deep (bottom) relative to the plane of initial diagnostic sectioning. (B) Snapshots of H & E-stained sections cut from the same sample as (B), after synchrotron imaging. Sections separated by roughly 25 micrometers are shown from most superficial to most deep, relative to the approximate plane of the pre-PBCT initial diagnostic sections.

### Histopathological Comparison of H&E and Micro-CT Imaging in Prostate Needle Biopsies

#### Case 1 (Benign)

Standard H&E histology demonstrated benign prostatic tissue, with classic undulating and tortuous prostatic glands with a small degree of partial atrophy. Cytoplasm was pale and eosinophilic. Nuclei were small without prominent nucleoli. Blood vessels of varying caliber were present within normal fibromuscular stroma. On micro-CT imaging, glands were clearly distinguishable from background stroma. The cytoplasm appeared pale with minimal density, while epithelial cell nuclei appeared dark and stood out against the background, proving to be the most useful feature for appreciating gland morphology. Glands appeared in well-circumscribed groups with non-infiltrative architecture, an observation that was further emphasized when scrolling through different levels of depth. The fibromuscular stroma appeared as dense fibromuscular bundles admixed with scant, less dense loose connective tissue, just as observed in standard H&E sections. Stromal cell nuclei did not stand out prominently and were not readily distinguishable from tumor nuclei (Fig 2). Blood vessels appeared as empty spaces with recognizable vascular architecture, though endothelial cell nuclei were difficult to discern. We are able to examine questionable cribriform architecture in deeper sections using micro-CT (Figure 3). Scrolling through the micro-CT stack revealed this to be benign with tufted cells in contact with one another, which was also apparent on serial H&E sections (Figs 3, S5).

#### Case 2 (Prostatic Adenocarcinoma, Gleason Score 3+3=6)

H&E examination revealed usual acinar type prostatic adenocarcinoma with crowded round glands, enlarged nuclei with prominent nucleoli, frequent intraluminal amorphous pink secretions, and infiltrative architecture among benign glands. On micro-CT, regions of prostatic adenocarcinoma demonstrated crowded round glands similar to H&E findings (Fig 4). Scrolling through the cluster revealed a minor degree of gland branching but no fusion of adjacent glands. It was difficult to distinguish benign from malignant glands in areas with numerous crowded glands on a single image. Micro-CT-enabled scrolling through these areas allowed us to study the infiltrative architecture of entire glands, including carcinoma and non-infiltrative benign glands (Figure S6). Nucleoli were not as apparent on micro-CT as in H&E sections. Flocculent material of variable density was present in gland lumina. Distinguishing atrophy from pattern 3 adenocarcinoma was more challenging in grey-scale micro-CT 5 micron MIPs compared with standard H&E-stained sections.

**Figure 4:**
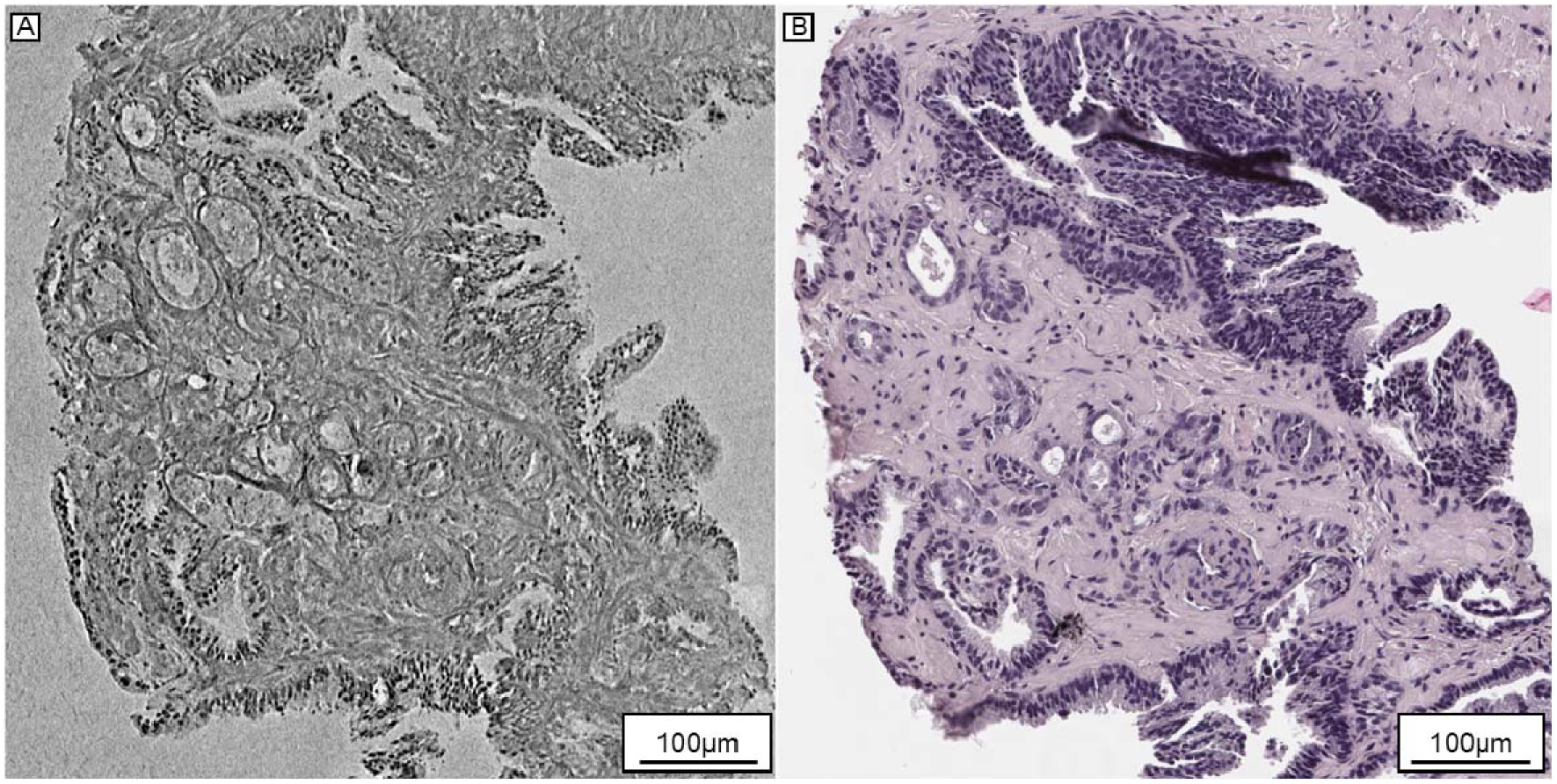
Morphology defining Gleason pattern 3 and benign partial atrophy visualized by PBCT and matching histology. (A) 8-slice maximum-intensity projection (∼4.8 micrometers thick) cropped from the reconstruction of case 3 to showcase morphology consistent with Gleason pattern 3. (B) H&E-stained histology extracted from case 3 after scanning, cropped to match the same approximate region as (A). Crowded round glands consistent with pattern 3 adenocarcinoma are visible across each modality, but regions of atrophy (lower right acini) were more difficult to discriminate within micro-CT than the histology.

#### Case 3 (Prostatic Adenocarcinoma, Gleason Score 4+4=8)

H&E histology demonstrated classic pattern 4 cribriform-type adenocarcinoma. A portion of the tumor exhibited less classic cribriform architecture with more vacuolar-appearing spaces, and another portion appeared as poorly-formed glands. Tumor cells had large nuclei with frequent prominent nucleoli. On micro-CT, the cribriform structure was readily identified, with the same architecture as seen on H&E (Figure 5). Similar to Case 2, areas that were easily distinguishable as benign glands surrounded by carcinoma on H&E were much more difficult to differentiate on single slice micro-CT, which is thinner than the amount of tissue represented in H&E sections. Scrolling through glandular z-stacks clearly showed benign glands connecting to more clearly benign glands and cancer connecting to more obvious cancer.

**Figure 5:**
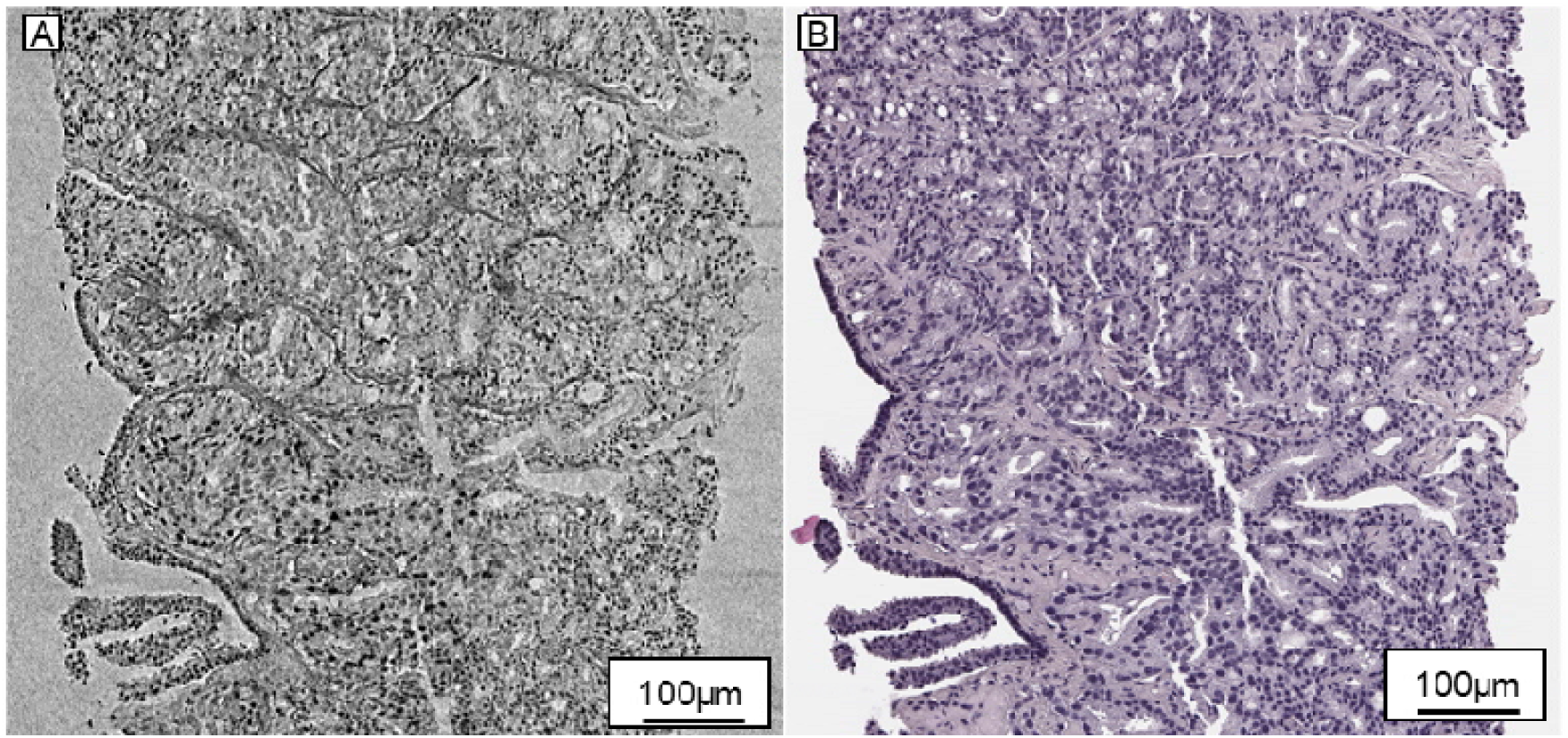
Gleason score 4+4=8 with cribriform architecture revealed by PBCT and correlative histology. (A) PBCT ROIs cropped from the 3D reconstruction of a core with a Gleason score of 4+4, resliced to qualitatively match the plane of histological sectioning. 3 8-slice maximum intensity projections (MIPs), corresponding to ∼4.8 micrometer virtual “slabs” (B) Correlative histology taken from the same core as (A) after synchrotron imaging. ROIs approximately matching the same tissue as (A) are shown as visualized by serially sectioned 5-micrometer H&E slides separated by approximately 25 micrometers of tissue depth.

#### Case 4 (Prostatic Adenocarcinoma, Gleason Score 4+5=9)

H&E slides demonstrated prostatic adenocarcinoma, predominantly cribriform pattern 4, with regional comedonecrosis, interpreted as Gleason score 4+5=9. The architecture revealed by H&E sections suggested intraductal prostatic carcinoma, though this was not confirmed by immunohistochemistry. Tumor cells exhibited large nuclei with prominent nucleoli. Much of the tumor had fragmented during processing, with fragments of cancer within empty spaces that were loose and disconnected, along with necrotic fragments. Part of the biopsy core showed reactive fibrosis and hemosiderin-laden macrophages (Figure S7). A focus of calcium phosphate was present adjacent to the cancer (Fig 6). On micro-CT, the same pattern 4 cribriform architecture was evident. Case 4 evaluation was more difficult than Case 3 due to fragmentation. The necrosis appeared as amorphous material of medium density (Figure 6). Nucleoli were not as apparent on micro-CT. The region containing hemosiderin-laden macrophages appeared as a black smear, consistent with high-density material (Figures 6, S7).

**Figure 6.**
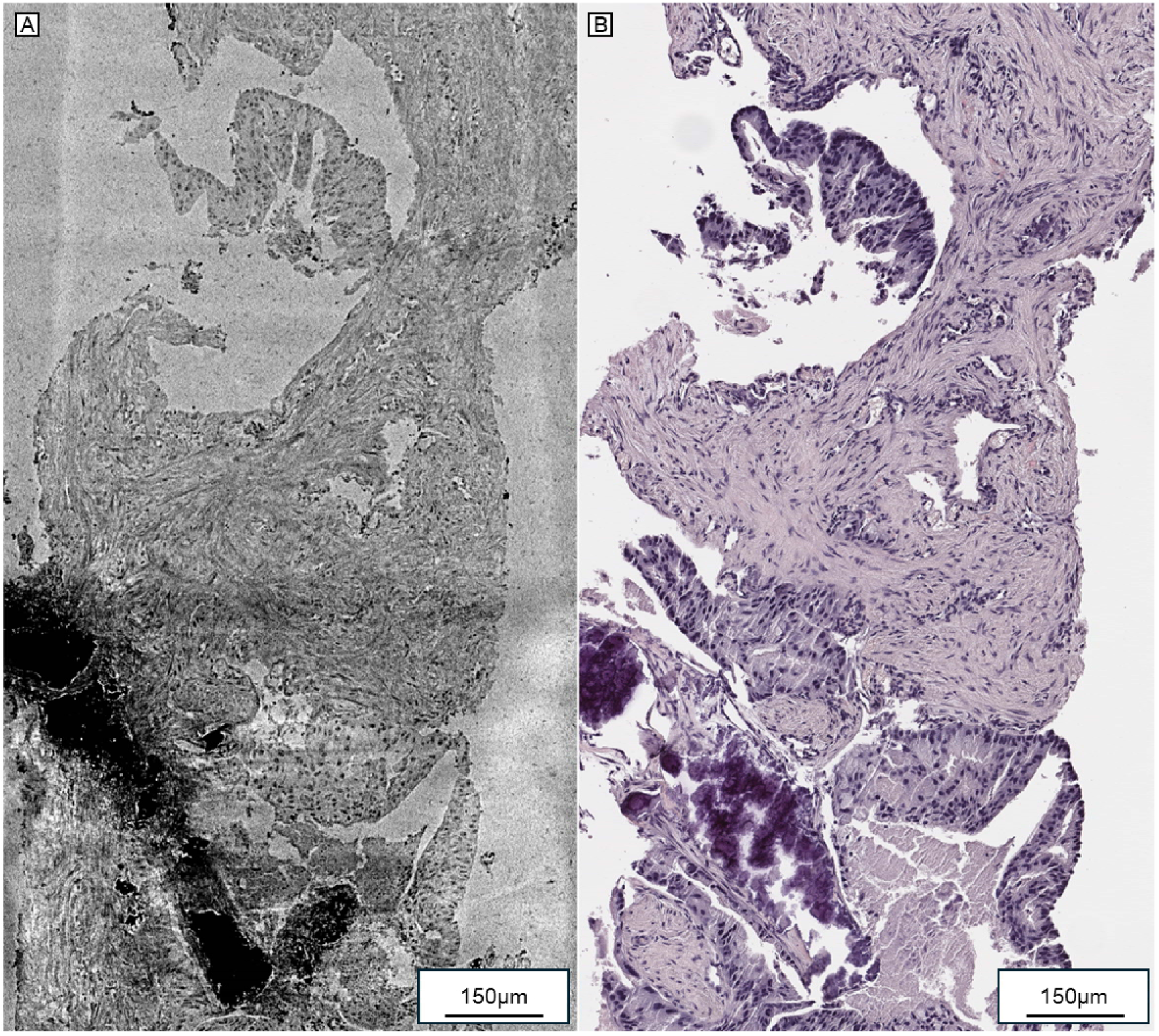
Gleason score 4+5=9 with comedonecrosis and incidental calcium phosphate crystals. (A) 8-slice inverted maximum intensity projection (∼4.8 micrometers thick) cropped from a 3D reconstruction of sample 4, which was graded as Gleason 4+5=9. A calcium phosphate deposit is visible as a large black patch of high-intensity pixels, but its internal structure is visible with contrast adjustment of the image (see data availability for Case 4). (B) Correlative histology of the matching tissue section collected from sample 4 after PBCT scanning. Comedonecrosis is present at the bottom right, adjacent to the calcium phosphate.

### Micro-CT facilitates segmentation of nuclei and virtual H&E staining

To automate the segmentation of nuclei clearly visible in our phase-based micro-CT reconstructions, we applied semi-supervised learning. PBCT enhances contrast between materials of different densities. Coupled with increased photon attenuation at a lower beam energy (14 keV), nuclei appear most prominent within our biopsy reconstructions (Figures 1-6).

We trained a 3D U-Net model from StarDist with a sparse set of hand-annotated training data and generated 3D instance segmentation masks representing candidate nuclei (Figure S8). As a complementary proof of principle, we used Ilastik random-forest classification to separate glandular spaces from gland tissue and candidate nuclear structures (Figure S9). Here, instance segmentation refers to assigning each labeled candidate nucleus a unique identifier. Given the small sample size of our dataset, the goal of segmentation within this case study was not to generate an approach for de-novo diagnosis, but to demonstrate that plausible, micrometer-scale morphologic detail corresponding to histopathology could be extracted in 3D with PBCT. Results were reviewed qualitatively with study pathologists (GC, JW) and were assessed with H&E-stained sections from the same cores taken after 3D scanning (Figure S8), illustrating compatibility with publicly available deep learning models.

To assess the feasibility of virtual H&E staining with our label-free micro-CT images, we asked whether a cycleGAN could produce plausible results after limited training. We selected representative 2D snapshots from the micro-CT reconstructions of Cases 1 and 3 and paired them with tissue-matched ground truth H&E images for training (see Methods). Training required <24 hours on a laptop RTX 4090 GPU, with model stabilization in <200 epochs. We observed that cycleGAN output strongly matched the texture, color, and microscale tissue architecture of ground-truth H&E (Figure 7, Figure S10). Images were reviewed with genitourinary pathologists and found to exhibit few features that were unsupported by the input image within our cohort. At low image magnifications, we observed output from our sparsely trained model to be difficult to distinguish from ground-truth images. Fine-tuning of simulations and quantitative validation of virtual staining fidelity and segmentation accuracy across a statistically powered cohort, while beyond the scope of this proof-of-concept study, represents an important direction for future work.

**Figure 7:**
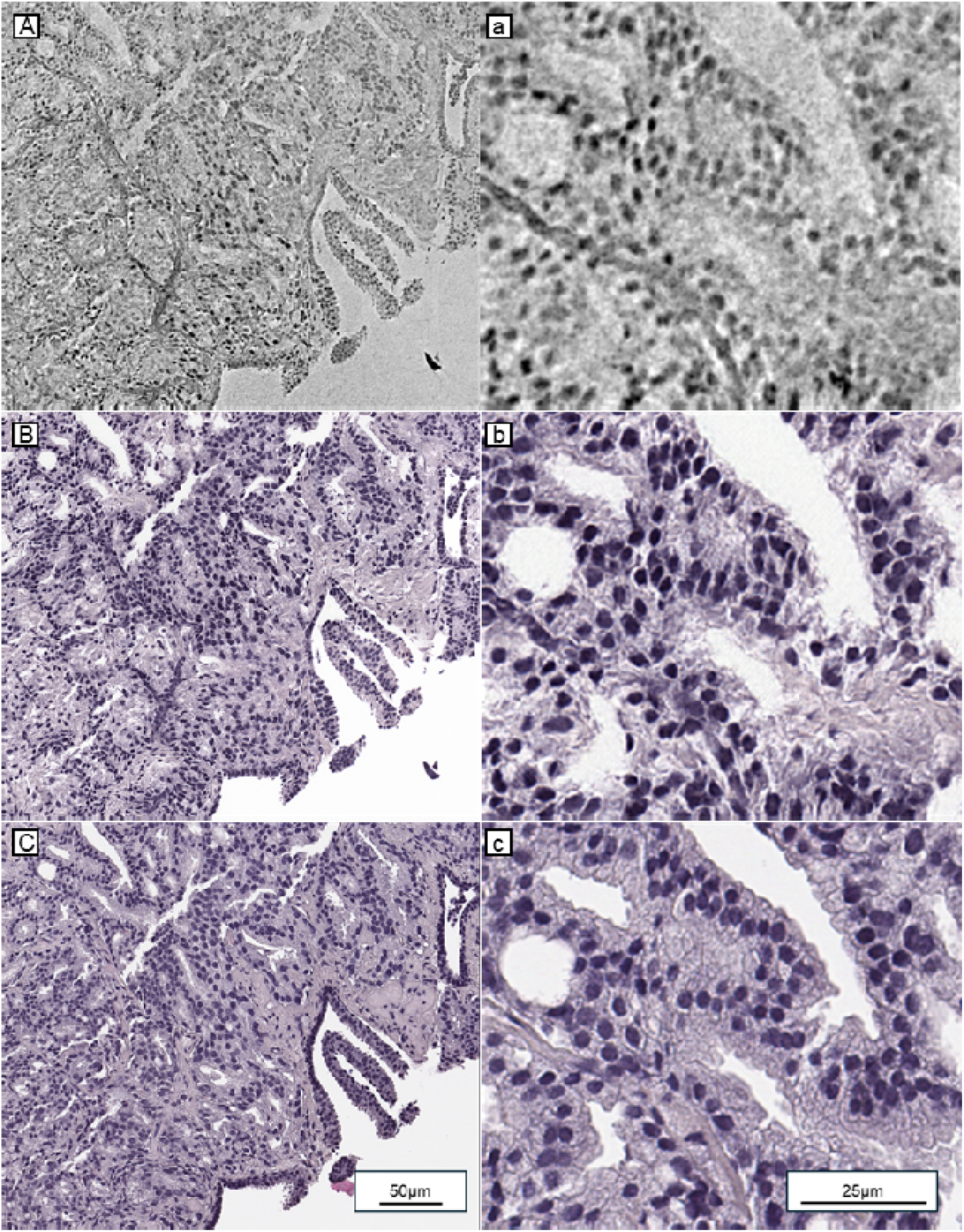
PBCT with tissue matched histology enables virtual staining. (A) 8-slice maximum intensity projection from the PBCT reconstruction of Case 3, with a higher-powered view of the same tissue shown in (a). (B, b) CycleGAN virtual staining of A and a respectively. (C) Conventional H&E-stained histology slide taken from Case 3 after PBCT scanning. The digitally-scanned slide was computationally aligned to the tissue shown in (A) to facilitate comparison.

## Discussion

Cancer tissue diagnosis based on traditional histopathology by nature is accomplished using sparse sampling of 3D tissue and lacks quantitative volumetric characterization of cells and tissues (Cheng et al., 2022; Ding et al., 2019; Liu et al., 2021). The desire for improved volumetric sampling has driven work towards 3D tissue imaging, which requires both sufficient resolution and field-of-view (FOV) to achieve equivalent or superior histopathological diagnostic power. Here we have explored the potential of undistorted multiangle visualization using the isotropic 3d images (with cubic rather than distorted rectangular voxels) of X-ray microtomography (histotomography, Ding et al. 2019). Histotomography was originally based on metal staining after fixation, which resulted in images that are based on both absorption and phase effects, dominated by the former. The problem addressed here was how to potentially utilize the power of histotomography with minimal change of existing clinical workflows that involve fixation and embedding in paraffin followed by histology. Phase contrast based on normal variation in tissue density reveals nuclear-scale detail in 3D cancer tissue volumes without the distortions characteristic of tissue sectioning.

Prostate cancer patients suffer from over and under treatment due to limitations in diagnosis and grading, which has driven work in 3D virtual histology that has shown promise for both basic research and clinically predictive value (Serafin et al., 2023; Song et al., 2024; Xie et al., 2022). In the present proof-of-concept study, we show that propagation-based phase-contrast micro-CT (PBCT), or phase histotomography, enables a 3D virtual histology starting with unstained FFPE human prostate needle-core biopsies. Resolution and contrast were sufficient for volumetric exploration of glandular and nuclear architecture at the micron scale, allowing one to distinguish between clinically significant phenotypes such as cribriform pattern 4 (Figures 1, 5) and a benign cribriform-like focus (Fig. 3). Here, we imaged <1 mm wide prostatic needle biopsies, but notably our imaging system cameras can image samples as wide as 5 millimeters with 0.61-micron voxel size (Sugarman and Vanselow et al., 2026; Yakovlev et al., 2022).

We have shown that phase-based micro-CT, or phase histotomography, allows one to distinguish between benign and malignant glands, to assign Gleason patterns, and to create serial sections in any region of imaged tumors in any plane of section. To facilitate histopathologic interpretation, cycleGAN based virtual H&E images can be simulated from the reconstructions. Post-PBCT serial section histology yielded paired training data mapping grayscale data to ground truth histology with sub-micrometer detail. At low to medium power, our cycleGAN inference images are nearly indistinguishable from ground truth histology (Figures 7, S10). Further work may increase the quality of high-power simulations. Our false-coloring results within a small cohort demonstrate the power of PBCT and correlative histology as training data for image-to-image AI models (Esposito et al., 2025; Zhang et al., 2025; Zhu et al., 2017).

Our results broaden the scope of the emerging field of 3D virtual histology, complementing the recent advances of light-sheet methods. Open-top light-sheet microscopy (OTLS) was used to measure 3D features for the risk stratification of prostate biopsies, enabling the use of AI models for the computational processing of their large datasets (Barner et al., 2023; Serafin et al., 2023; Song et al., 2024; Xie et al., 2022). However, fluorescence-based approaches to 3D virtual histology typically require deparaffinization, tissue clearing, and the addition of antibodies or other contrast enhancing stains - labor-intensive experimental steps outside of the workflow of pathology. Furthermore, these techniques are constrained by the penetration of visible light, which can be disrupted by calcifications (Figure 6), pigmentation, or especially dense/thick tissue samples.

Propagation-based phase contrast micro-CT has been shown to produce 3D images of other cancers without deparaffinization or the addition of stain. Topperwien and Salditt have characterized the use of PBCT and X-ray tomography in general for the high-resolution imaging of unstained soft-tissue biopsies in several different embedding media, including ethanol and paraffin wax (Frost et al., 2023; Töpperwien et al., 2018, 2019, 2020). Pinkert-Leetsch used synchrotron PBCT for the virtual histology of pancreatic cancer, segmenting the 3D shape of a pancreatic duct and identifying perineural invasion (Pinkert-Leetsch et al., 2023).

Here, we used synchrotron PBCT to generate 3D reconstructions across prostate needle cores after initial diagnostic sectioning. Several biopsies were serially sectioned, and H&E stained after volumetric imaging to provide direction comparison between micro-CT and histological representation of nearly identical biological space. Within our micro-CT reconstructions, we identified cell nuclei, fibromuscular stroma, and both benign and malignant prostate acini/lumen. We further delineated hallmark features of benign, Gleason pattern 3, Gleason pattern 4 (including cribriform/fused glands), and comedonecrosis of Gleason pattern 5 within our unstained, paraffin-embedded 3D X-ray images. We corroborated these features with traditional histopathology images taken before and after PBCT and evaluated by board-certified GU pathologists (GC, JW) (Figures 2-6). Several samples were serially sectioned, which also enabled us to explore and validate depth-based variation of glandular structure (Figures 2 and Figure S5). At present, 3D volumetric false coloring is not yet feasible due to sparse paired training data and distortions in correlative histology images associated with cutting, staining, and placement on glass slides. The results demonstrated here with sparse data encourage a strong prognosis for future studies, which could potentially incorporate histology images intentionally sectioned at different angles. The isotropic nature of PBCT reconstructions enables virtual re-slicing at any angle, facilitating image selection for a 3D model. Although nucleoli were more difficult to visualize within our micro-CT reconstructions, cycleGAN virtual staining appeared to produce images with nucleolus-like textures. Quantitative matching of nucleoli from H&E images and matching regions in PBCT scans across a larger dataset may yield improvements in virtual staining.

This line of research will benefit from micro-CT evaluation of additional Gleason patterns including mucinous fibroplasia/collagenous micronodules, poorly formed glands, glomerulations, solid sheets of tumor, and single cells, as well as additional benign findings, such as diverse examples of atrophy, basal cell hyperplasia, and granulomatous prostatitis. Rigorous AI-based segmentation of diagnostic features with histology cross validation would benefit from a study of biopsies without prior sectioning - either collected prospectively or synthetically from radical prostatectomy sections (as performed in Song and Serafin et. al) (Serafin et al., 2023; Song et al., 2024). While this would forgo strict comparison with an initial clinical diagnosis of biopsy cores, segmentation would avoid the image artifact associated with cut paraffin at the face of the sample (Table S1, Neuroglancer) (Maitin-Shepard et al., 2021). Our approach focuses on images attainable at a synchrotron (Advanced Light Source at LBNL), which users must apply to compete for limited time, making large-scale clinical studies difficult. However, benchtop micro-CT approaches have demonstrated single micrometer spatial resolution, and the 5 mm detector system employed in this study is usable in a benchtop setting (Yakovlev et al., 2022). Recent studies by Katsamenis, Eckermann, and others suggest the feasibility of the virtual histology of human tissues in a benchtop setting (Eckermann et al., 2020; Katsamenis et al., 2019; Töpperwien et al., 2018). As technology improves, larger scale studies of human cancer with sub-micrometer and better resolution will become tractable. While we did not train AI models on a large scale, our results suggest that PBCT makes plausible the segmentation of nuclei and lumen using pre-existing models with small amounts of training data (Figures S8, S9, Video S1). While further work involving a larger number of biopsies will be necessary to improve accuracy and feasibility clinical applications. PBCT can be used to study tumor heterogeneity, and to potentially add value to the practice of pathology.

## Author Contributions

KCC is responsible for the long-term vision of using X-rays to image cancer and other tissue in 3D and identified prostate cancer as an ideal proof of principle. PLR advised propagation-based phase contrast parameters, phase retrieval, and image reconstruction. JW precisely defined the clinical scope of this project, selected the cases, directed the IRB work with GC, and completed the manuscript with ALS and KCC. ALS suggested the use of phase rather than metal-stained micro-CT, prepared samples from paraffin blocks, performed synchrotron micro-CT imaging, reconstruction, image segmentation, virtual staining, coordinated work between KCC and ALS labs, and wrote the manuscript. DJV supported synchrotron imaging and reconstruction, developed the Neuroglancer and web visualization tools for data sharing, and performed image registration. EC and DP supported synchrotron imaging and image analysis at LBNL. JS oversaw conception of statistical aspects of the project, advised image segmentation and cycleGAN methods, and participated in manuscript writing. GC and JW performed clinical analysis of micro-CT and correlative histology data. All authors reviewed and edited the manuscript.

## Acknowledgements

We thank Jean Copper and Marianne Klinger for tissue embedding advice and histological sectioning of micro-CT imaged prostate, and Maksim Yakovlev, Carolyn Zaino, David Northover, Jessica Christ, and KC Ang for help with micro-CT imaging at Department of Energy synchrotrons at Argonne and Lawrence Berkeley National Laboratories. We acknowledge computational support provided by Penn State Institute of Computational and Data Sciences Roar Core Facility (*RRID: SCR*26424) and Pennsylvania State University RISE Core Facility (*RRID: SCR*26426). Additional research support provided by Penn State Institute for Computational and Data Sciences (*RRID: SCR*25154) and Penn State Center for Applications of Artificial Intelligence and Machine Learning to Industry (*RRID: SCR*22867). ALS was partially supported by 2025/2026 Rising Researcher Grant *ICDS RR*2527745 from Penn State’s Institute for Computational & Data Sciences (*RRID: SCR*025154). Imaging work was made possible by funding from NIH grants 1R24OD18559, R24OD035407 and from the Jake Gittlen Laboratories for Cancer Research and Penn State Cancer Institute and to KCC. This research used resources of the Advanced Light Source, which is a DOE Office of Science User Facility under contract no. DE-AC02-05CH11231. We acknowledge involvement of Zebrafish Functional Genomics Facility (RRID:SCR_021199) in the development of histotomography and instrumentation.

## Data availability

Interactive web visualization of the 4 main cases and their corresponding post-PBCT histology can be accessed with the links in Table S3. All other data and code reported in this manuscript are available upon request to the first, second, or corresponding authors.

### Ethics statement

All specimens were handled and investigated in accordance with IRB-approved protocols.

## Declaration of generative AI and AI-assisted technologies in the manuscript preparation process

During the preparation of this work the authors used OpenAI ChatGPT 5.x Pro for limited editorial assistance with author-drafted manuscript text. After using this tool/service, the authors reviewed and edited the content as needed and take full responsibility for the content of the published article.

## Supplement

**Table S1:** Tissue samples selected for PBCT.

| Case ID | Initial Diagnosis | Post-PBCT serial-sectioning (yes/no) |
| --- | --- | --- |
| 1 | benign | yes |
| 2 | 3+3 | yes |
| 3 | 4+4 | yes |
| 4 | 4+5 | yes |
| 5 | 3+3 | no |

**Table S2:** Tissue contrast key in histology and PBCT.

| Modality | Nuclei | Cytoplasm | Lumen | Primary source of signal |
| --- | --- | --- | --- | --- |
| Hematoxylin | blue/purple | minimal | white | Dye-mordant complex binds anionic moieties of DNA/RNA |
| Eosin | minimal | pink |  | Anionic dye binds cationic moieties of proteins |
| PBCT | white | light gray | gray | Phase changes ($\delta$) induced by variations in electron density; absorption ($\beta$) governed by Beer's law<br>$\delta/\beta \gg 1$ with hard X-rays ($>10\text{keV}$) |
| Inverted PBCT | dark | gray | light gray |  |

**Table S3:** Neuroglancer and histology links for sharing.

| Sample | Grade | Link |
| --- | --- | --- |
| 1 | benign | CT<br>https://cephalopod.team/histotomography/Octo9/Neuroglancer/?j=JRW004_invert_z31Histo_match_v1.15_nocontrols<br>Histology<br>https://chenglab.io/osd/daphioMirror/anatomy/histology/?t=JRW004&z=31&c=0.2824365653992985,0.3848448767286696,0.3110399999999999,0.19436481447963797 |
| 2 | 3+3 | CT<br>https://cephalopod.team/histotomography/Octo9/Neuroglancer/?j=JRW005_invert_z28Histo_match_v1.15_nocontrols<br>Histology<br>https://chenglab.io/osd/daphioMirror/anatomy/histology/?t=JRW005&z=28&c=0.5917754039605937,1.3961940698475308,0.2705428066888435,0.1674178235027779 |
| 3 | 4+4 | CT<br>https://cephalopod.team/histotomography/Octo9/Neuroglancer/?j=JRW006_invert_z10Histo_match_v1.15_nocontrols<br>Histology<br>https://chenglab.io/osd/daphioMirror/anatomy/histology/?t=JRW006&z=10&c=-0.023479754009266385,0.04277554734193584,1.1748974102883647,0.8198543140092132 |
| 4 | 4+5 | CT<br>https://cephalopod.team/histotomography/Octo9/Neuroglancer/?j=JRW007_invert_z35Histo_match_v1.15_nocontrols<br>Histology<br>https://chenglab.io/osd/daphioMirror/anatomy/histology/?t=JRW007&z=35&c=-0.10899380611053566,0.11971650827109381,1.2740068917474268,1.8945361105726821 |

**Figure S1:**
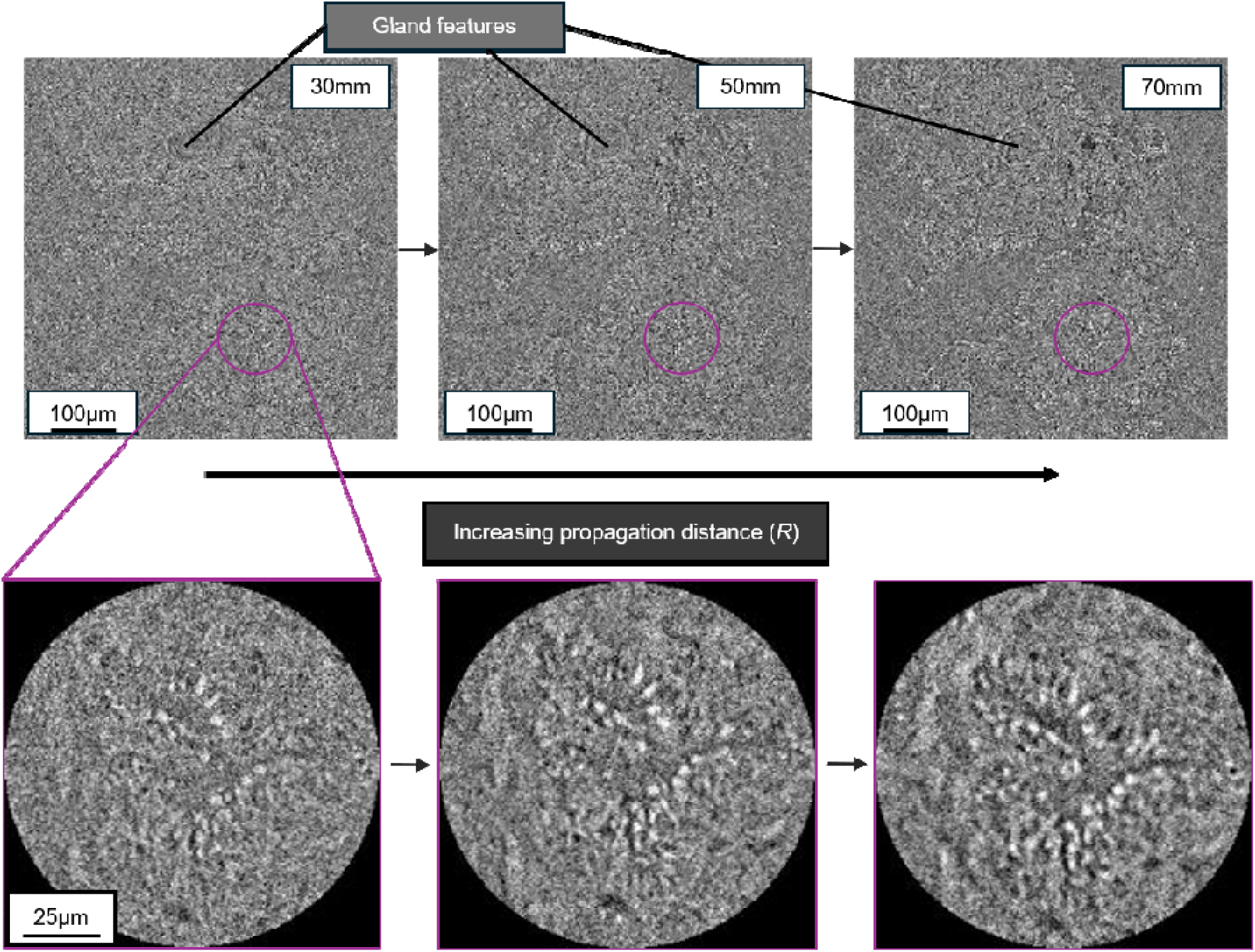
Propagation-based edge enhancement at 30mm, 50mm, and 70mm of propagation distance with 14keV parallel beam X-rays without phase retrieval. (Left to right) 2D slice of the same 3D PBCT scan shown at increasing sample-to-scintilla or (propagation) distance. Reconstructions shown here were performed without the use of phase retrieval or a bilateral filter to preserve phase artifacts for comparison. Glandular features and cell nuclei are readily visible at each distance but show significantly improved contrast at 50 and 70mm as compared to 30mm. An example of cell nuclei surrounding glandular lumina are labeled with a purple circle in each snapshot.

**Figure S2:**
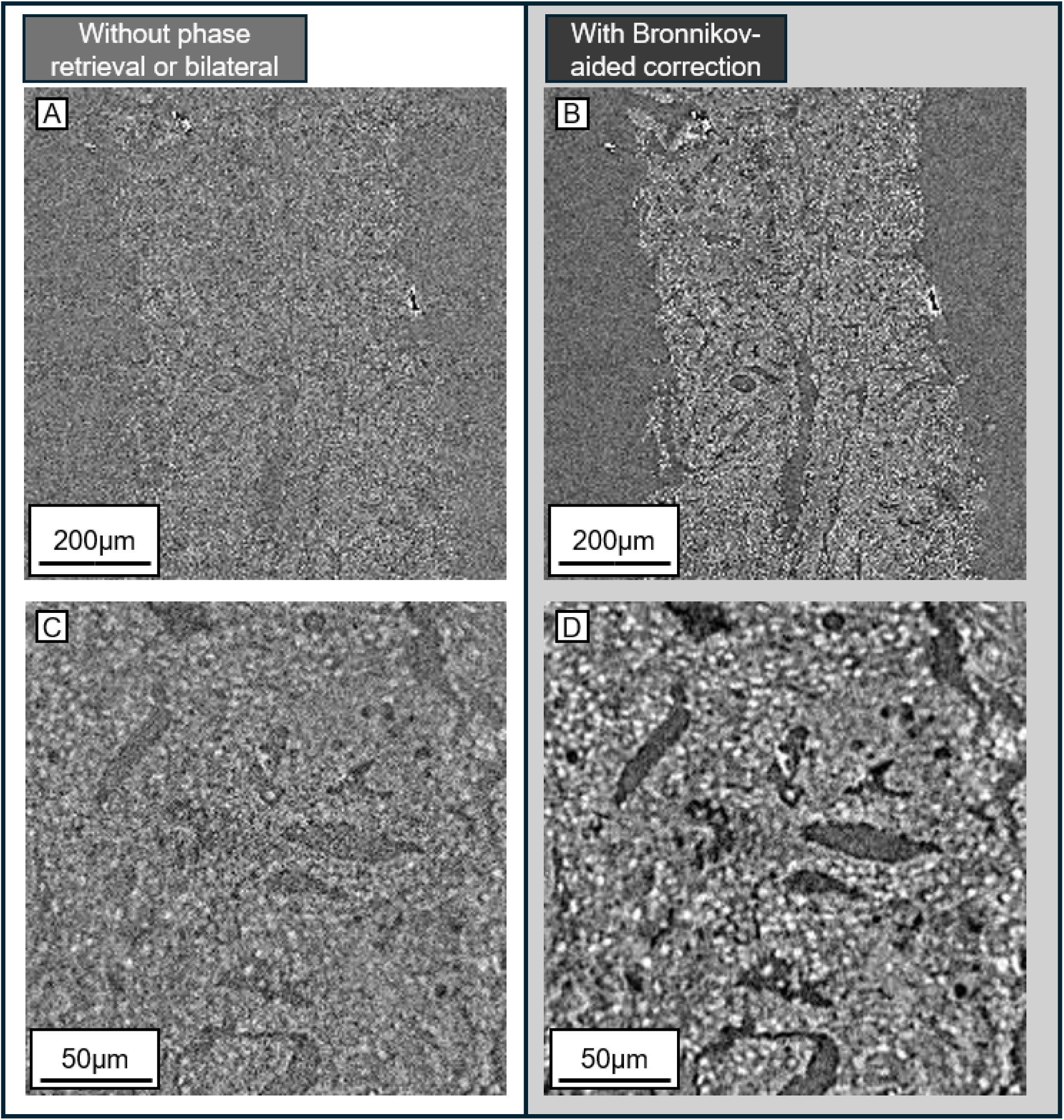
Phase retrieval with Bronnikov-aided correction (BAC) improves contrast while preserving nuclear-scale features. (A) 2D slice of a Gleason pattern 4+4 ROI taken from a scan performed at 50mm propagation distance. Reconstruction of this slice was performed without phase retrieval or bilateral filter. (B) The same ROI from the same sample as (A) reconstructed with the Bronnikov-aided correction (parameter values 2 and 1.25, package = HoToPy) for phase retrieval. (C) Higher zoom snapshot taken from the same reconstruction as (A). (D) Higher zoom snapshot from the same reconstruction as (B)

**Figure S3:**
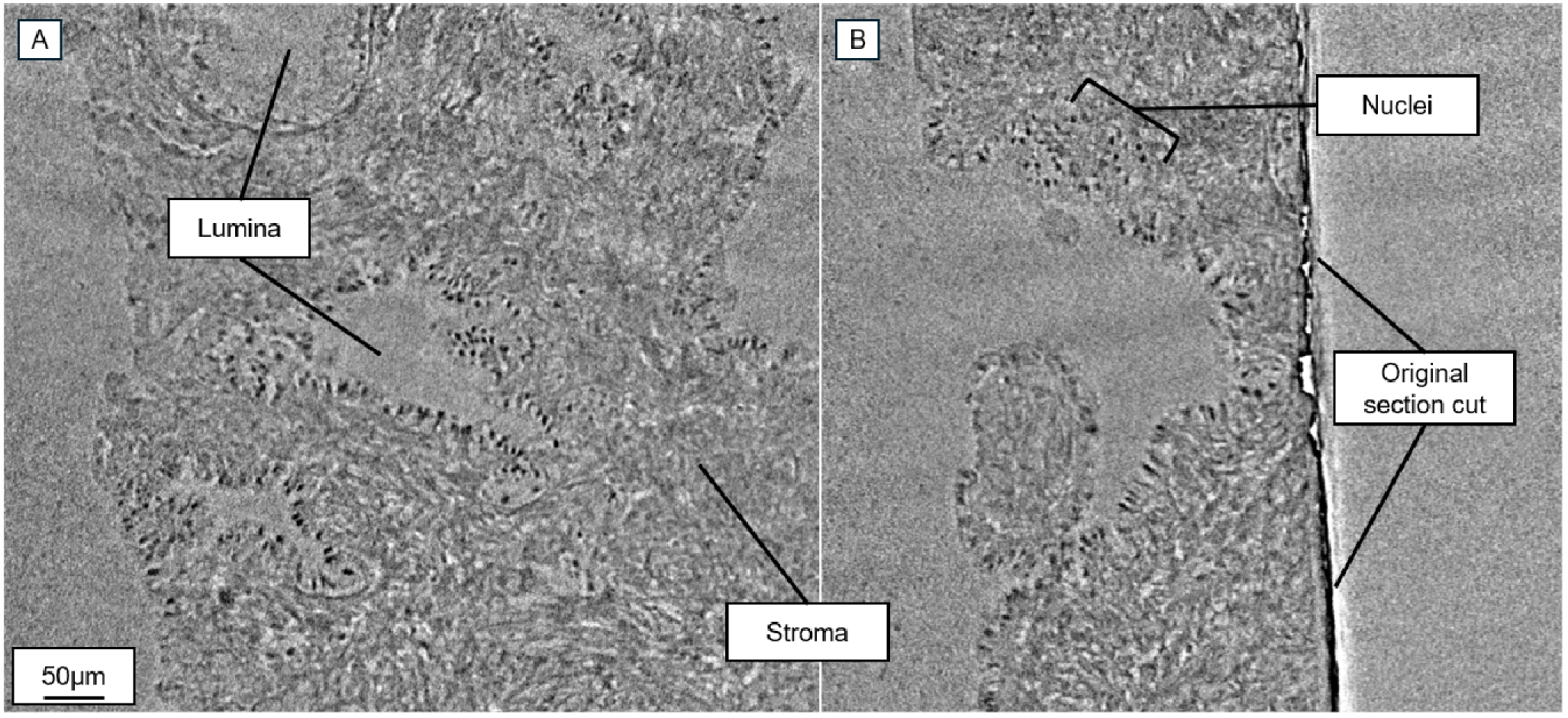
Isotropic resolution of PBCT enables visualization of Gleason grade-defining features in any plane. (A and B) Single (0.65-micrometer depth) orthogonal slices taken from a reconstruction of Case 5. (A) PBCT reconstruction resliced to qualitatively match the plane of initial histologic sectioning. (B) Orthogonal view with respect to (A), highlighting the paraffin edge induced by the microtome and the structures beneath the surface, including nuclear detail.

**Figure S4:**
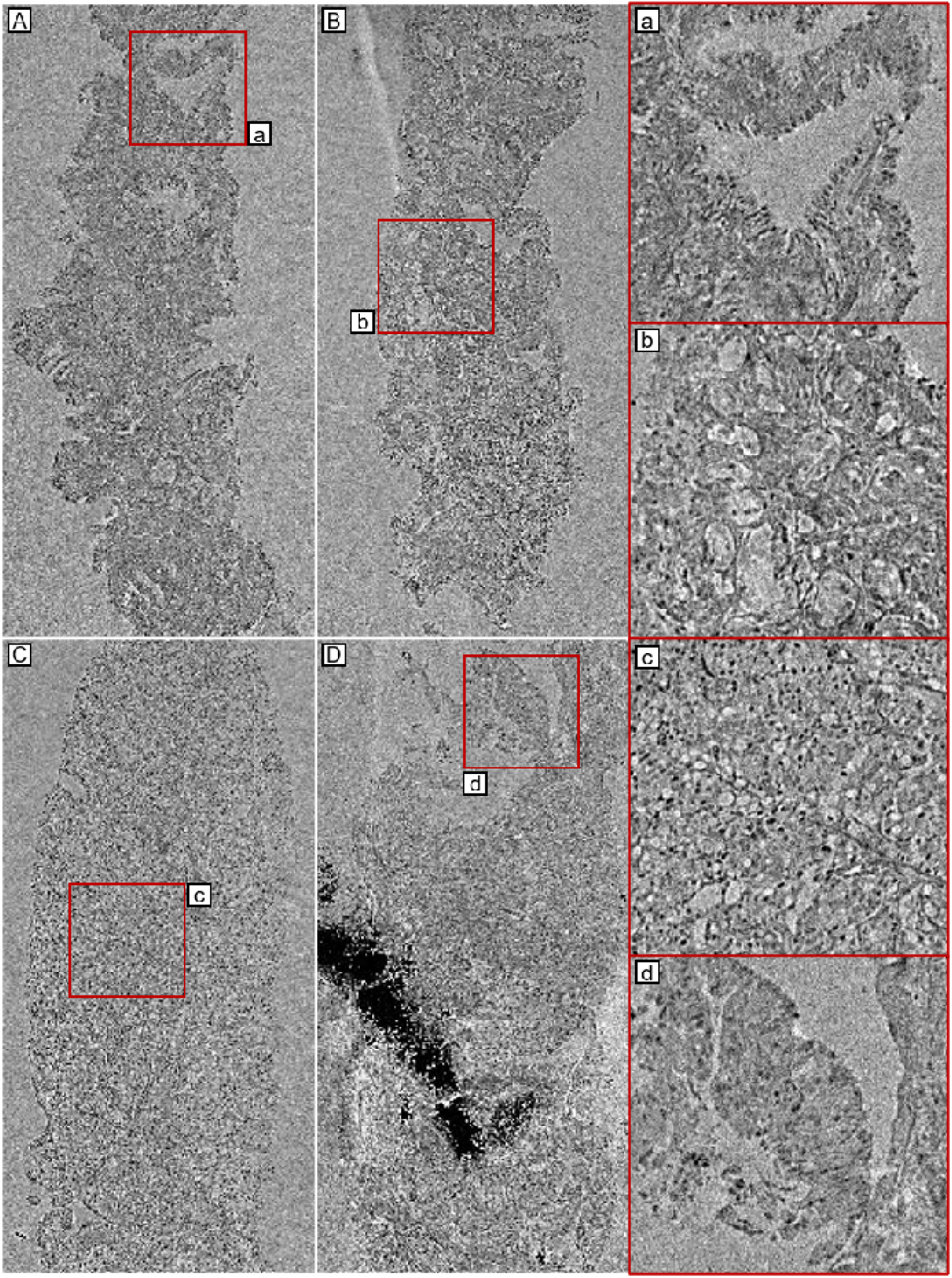
PBCT resolves Gleason grade-defining features from benign tissue to pattern 4+5. (In order A-D) Single virtual-slice ROIs from PBCT needle-core biopsy reconstructions with initial diagnoses of benign (Case 1), pattern 3+3 (Case 2), pattern 4+4 (Case 3), and pattern 4+5 (Case 4). The large patch of high attenuation (dark pixels) captured in D corresponds to calcium phosphate deposits. (a-d) Higher-powered insets selected from regions of each biopsy marked by a red square. (a) Benign gland, (b) cluster of pattern 3 glands, (c) cribriform pattern 4+4 glands, and (d) pattern 4+5 with necrosis.

**Figure S5:**
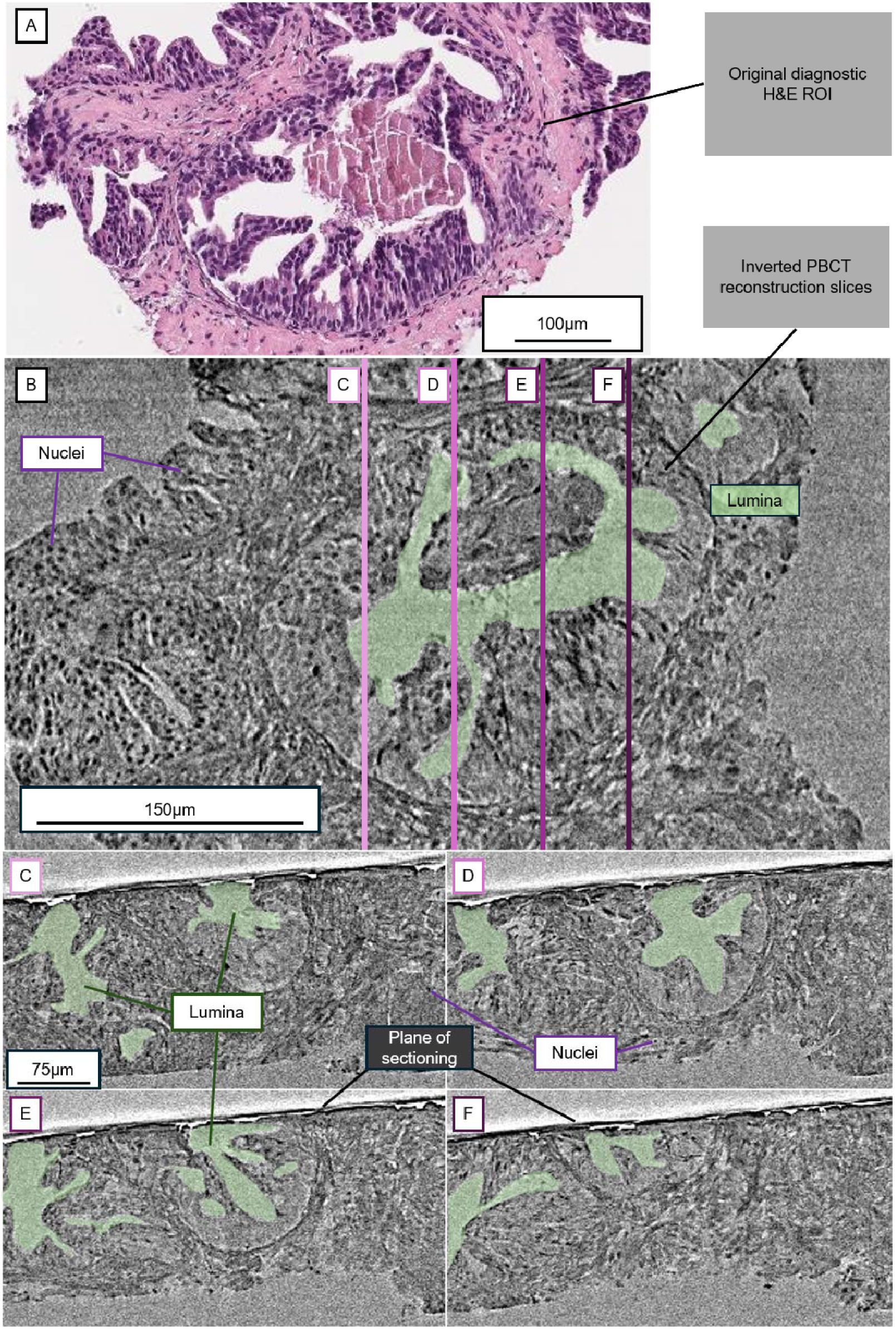
Orthogonal virtual sections reveal variation in prostate gland architecture beneath the plane of sectioning. (A) Virtual single-slice PBCT section of a benign needle-core, cropped to an ROI with features representing prostatic intraepithelial neoplasia (PIN). Glandular lumina and cell nuclei are denoted by green and purple labels, respectively. 4 orthogonal virtual sections separated by approximately 60 micrometers are shown through the gland, denoted by B, C, D, and E. (B) Orthogonal section to (A) illustrating the variation in luminal architecture below what is visible at the cut plane where initial diagnostic slides where sectioned. (C) Additional plane orthogonal to (A), parallel to (B), and ∼60 micrometers laterally spaced from (B) and (D). (D) Orthogonal virtual section to (A) ∼60 micrometers from (E) illustrating fused glands below the surface of the plane where the initial diagnostic section was cut for this core.

**Figure S6:**
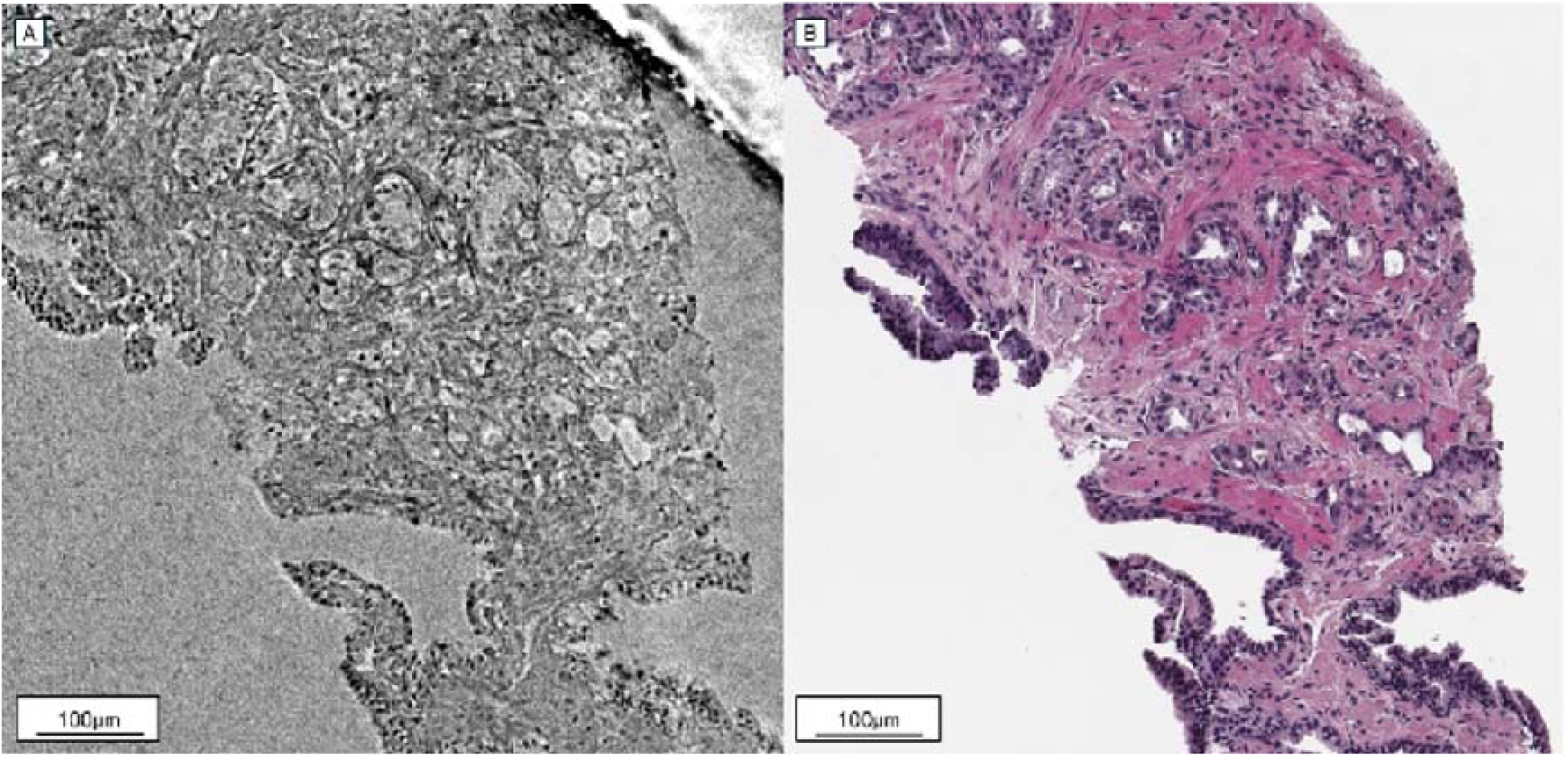
Infiltrative carcinoma visualized by 3D PBCT and serial-section H&E. (A) Single 0.61 micrometer-thick slice taken from a 3D reconstruction of prostate tissue classified as Gleason pattern 3 (Case 2). (B) Correlative H and E-stained histology of the same tissue region captured after PBCT scanning. Crowded round glands consistent with acinar type prostatic adenocarcinoma are visible. Luminal spaces serve as tissue landmarks, and crowded, poorly-formed acini consistent with infiltrative adenocarcinoma are visible across each modality.

**Figure S7:**
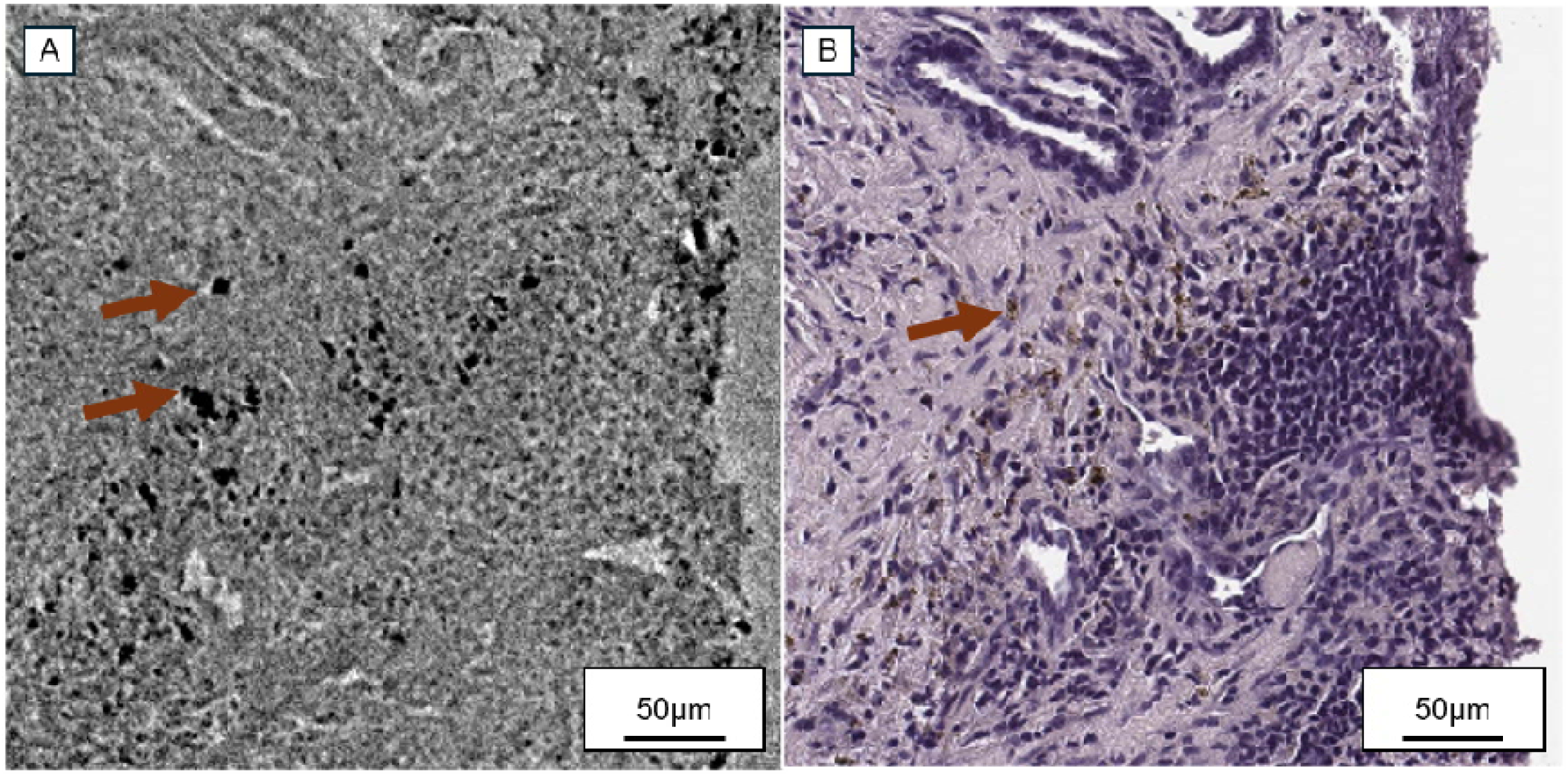
PBCT resolves attenuation consistent with hemosiderin-laden macrophages. (A) Single 0.61 μm-thick slice taken from the reconstruction of case 4, oriented to qualitatively match the plane of post-PBCT correlative histology. (B) Post-PBCT correlative H and E-stained histology of the tissue sample taken from case 4. Hemosiderin laden macrophages are visible as brown/rust-colored deposits (exemplar marked with an arrow).

**Figure S8:**
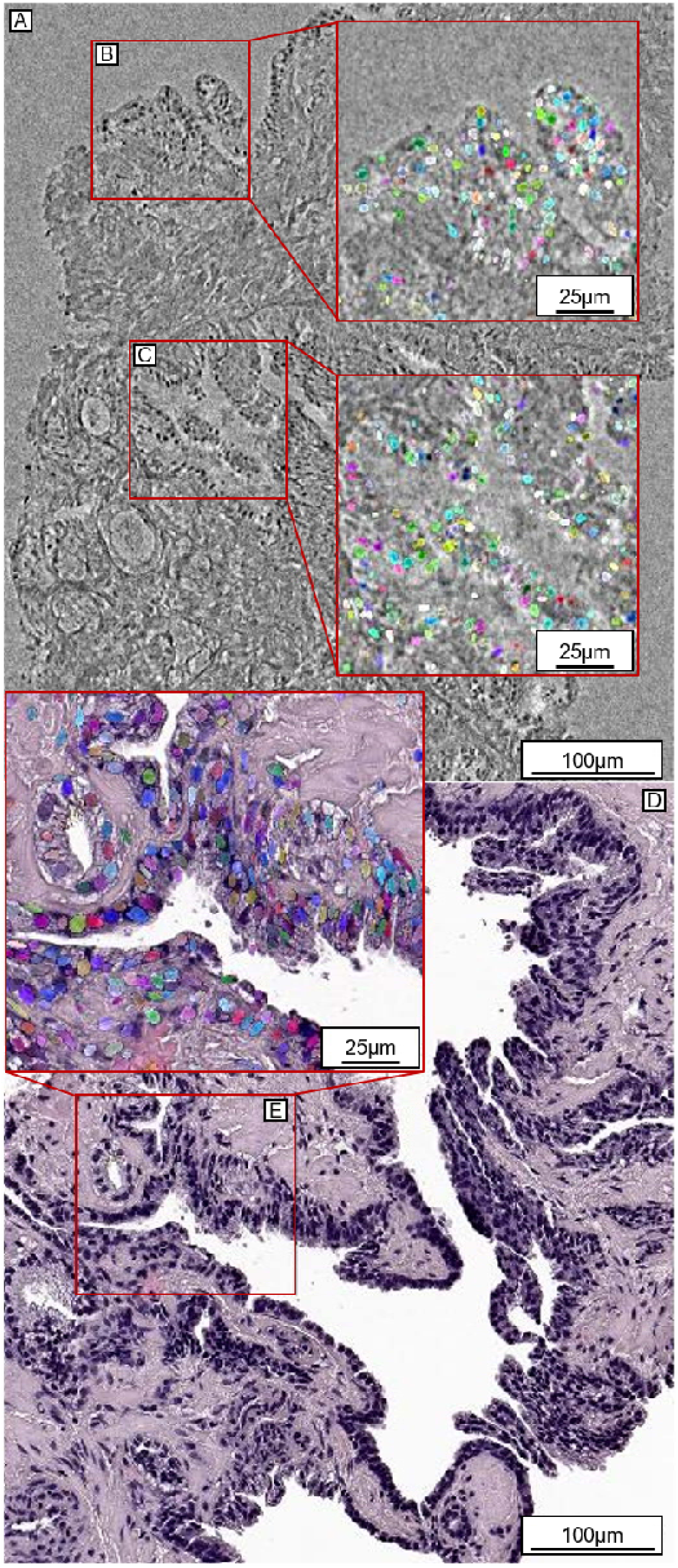
Segmentation of candidate nuclei in 3D PBCT and 2D histology with StarDist. (A) Single-slice ROI taken from a PBCT reconstruction of a pattern 4+4 core biopsy. (B) ROI from the boundary of the core with StarDist 3D U-Net segmentation results rendered as a multicolored instance segmentation. (C) ROI of a prostate gland with StarDist segmentations rendered as in (B). (D) Correlative histology taken from the same core as (A-C) after synchrotron imaging. (E) StarDist 2D U-Net segmentation results rendered as a multicolored instance segmentation as an example of preliminary nuclear detection in correlative histology.

**Figure S9.**
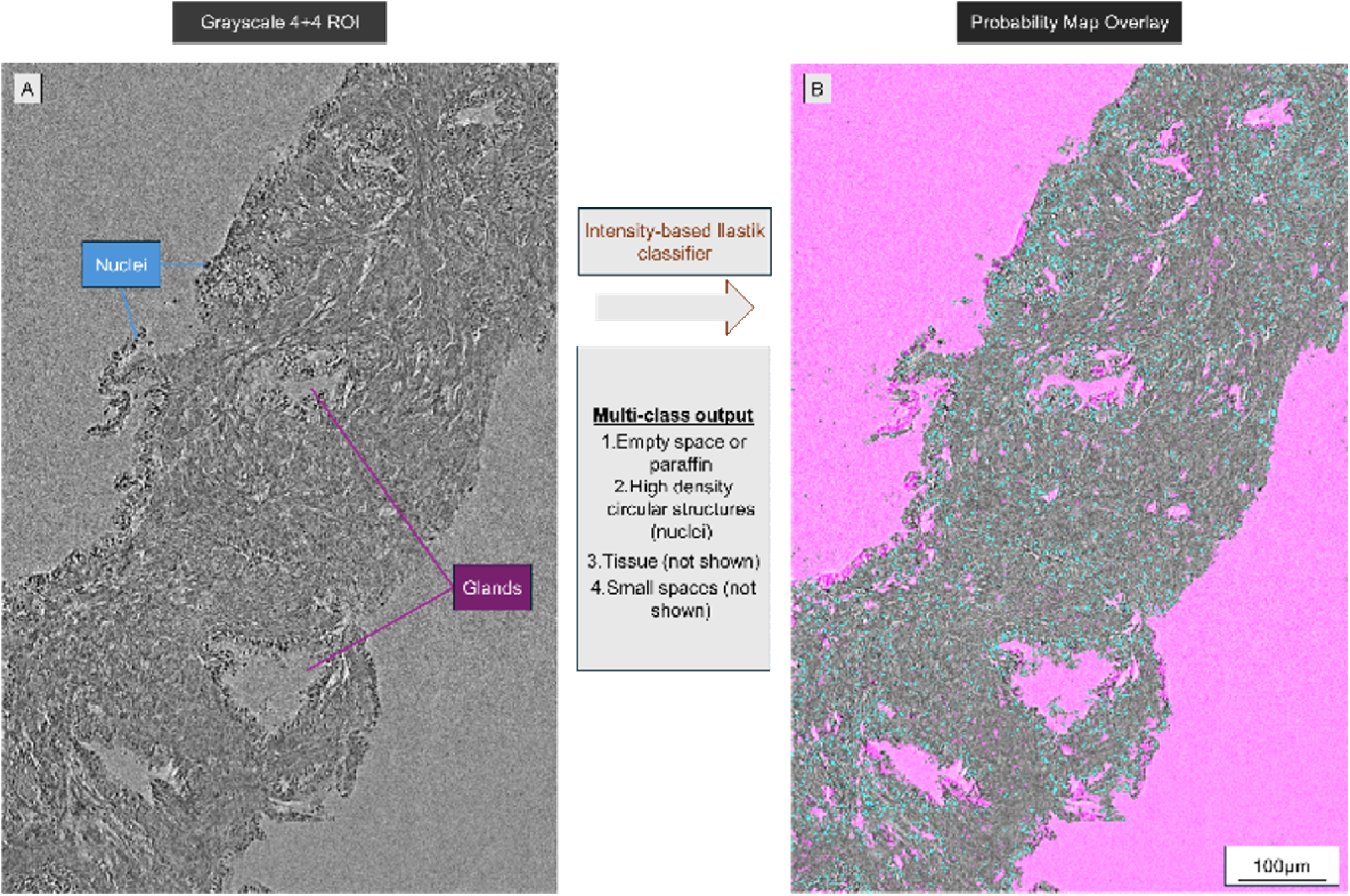
Ilastik random-forest classification segments gland space, tissue, and nuclear structures. (A) Inverted grayscale ROI taken from a Gleason score 4+4 (Case 3, ISUP grade 4) biopsy reconstruction. The 2D slice in (A) was resliced and cropped to mimic the plane of histological sectioning. Nuclei (blue arrow labels, visible as black dots) and glands (magenta arrows) are revealed by PBCT. (B) Ilastik segmentation result visualized as a probability map composite overlay in ImageJ. Nuclei are colored as blue/cyan while glandular lumen and background space are labeled as magenta.

**Figure S10:**
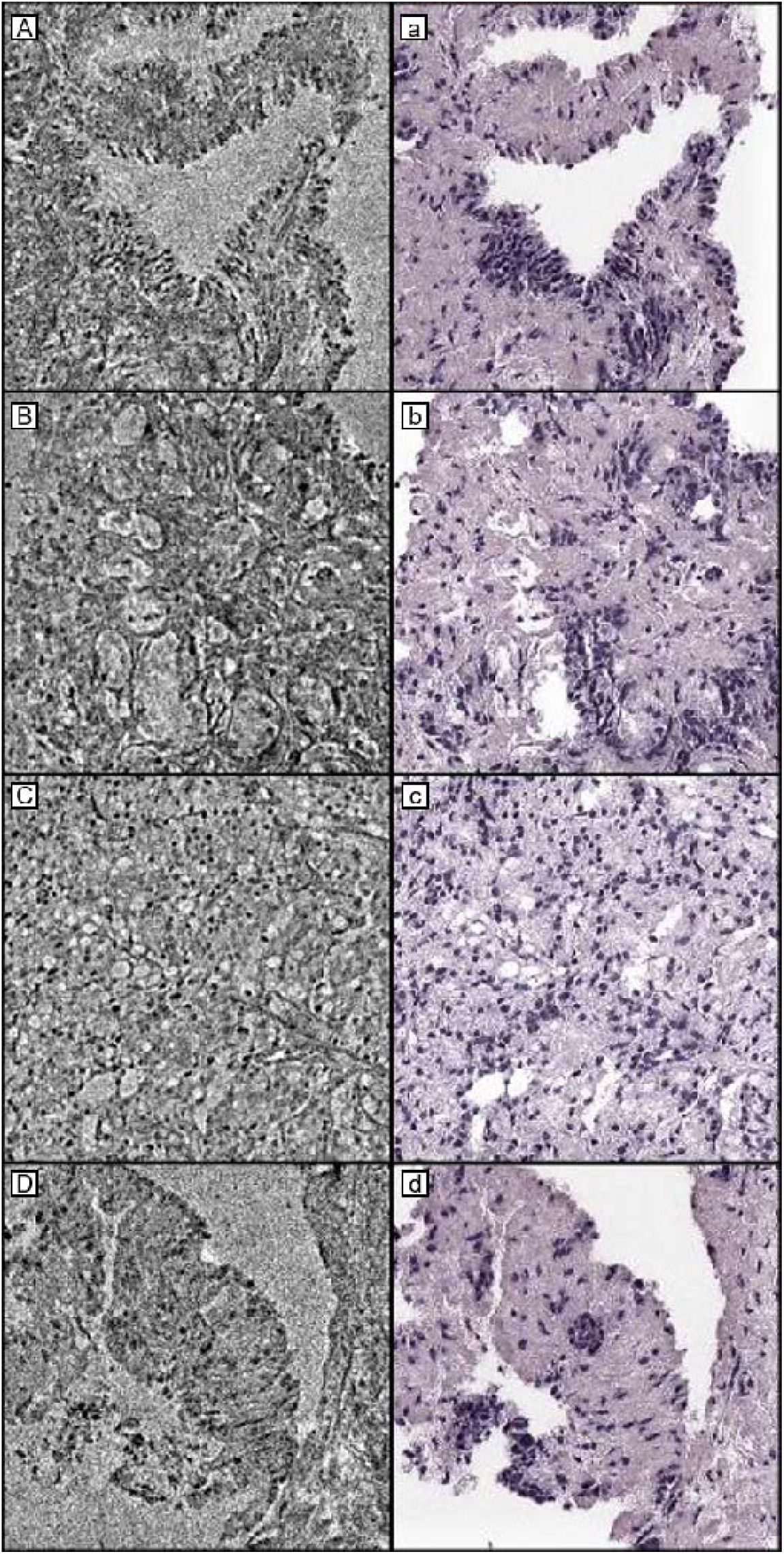
PBCT enables virtual staining. (In order A-D) Single virtual-slice ROIs from PBCT reconstructions of Case 1 (benign, A), Case 2 (3+3, B), Case 3 (4+4, C), and Case 4 (4+5, D). (a-d) Corresponding images virtually stained with a cycleGAN trained with snapshots from cases 1 and 3.

